# Sex-dependent cortico-amygdala circuits controlling emotion recognition

**DOI:** 10.1101/2025.10.28.685103

**Authors:** Jose Antonio González-Parra, Irene P. Ayuso-Jimeno, Xiao Chang, Paula Somoza-Cotillas, Vittoria Acciai, Marc Canela-Grimau, Laura Vidal-Palencia, Liwei Tan, Bowen Deng, Francesco Papaleo, Gunter Schumann, Arnau Busquets-Garcia

**Author notes:** Correspoding author: Arnau Busquets-Garcia, PhD, Cell-Type Mechanisms in Normal and Pathological Behavior Research Group Neuroscience Research Program, Hospital del Mar Research Institute. PRBB, Calle Dr. Aiguader 88 Barcelona 08003 SPAIN, Mail.

## Abstract

In social species, the ability to recognize others’ emotional states is essential for appropriate social interactions, yet it often declines with age and is impaired in various neurodevelopmental and neurodegenerative disorders. While emotion recognition has been characterized in both humans and rodents, the underlying neural circuits and how they vary by sex and age remain poorly understood. Here, we used a negative Emotional state Discrimination Task (EDT) in TRAP2 transgenic mice to map brain regions engaged during negative emotion recognition in young and aged animals. Young male and female mice successfully discriminated emotionally altered conspecifics, recruiting the basolateral amygdala (BLA) and medial orbitofrontal cortex (MO) in a sex-specific manner. Fiber photometry revealed distinct activation dynamics in these regions, and chemogenetic inhibition of bidirectional BLA-MO projections abolished emotion recognition in male but not female mice. Notably, young human participants also showed sex-specific recruitment of BLA and OFC during negative facial emotion recognition. Moreover, aging selectively impaired emotion recognition in male mice, coinciding with reduced BLA activity. Remarkably, chemogenetic activation of BLA in aged male mice rescued this deficit. Together, these findings identify a sex-dependent BLA-MO circuit as a conserved neural substrate for emotion recognition and demonstrate that age-related impairments can be reversed through targeted circuit-level intervention.

## Introduction

Social cognition encompasses the ability to perceive, interpret, and integrate socio-emotional information, while regulating behavior to facilitate adaptive social interactions among conspecifics^1,2^. Emotion recognition, which is a core component of social cognition, enables individuals to infer others’ emotional states through behavioral responses, facial expressions, vocalizations, and body language^3^. Impairments in emotion recognition are prevalent across neurodevelopmental^4^, neuropsychiatric^5^, and neurodegenerative disorders^6^. Moreover, biological variables such as sex^7,8^ and age^9^ influence emotion recognition abilities, yet the neural mechanisms underlying these effects remain poorly understood.

Recent evidence indicates that pro-social behavioral responses including emotion recognition and expression are not exclusive to humans but are also observed in other mammals such as rodents^10–14^. Thus, rodents discriminate emotional states of conspecifics^12,13^, exhibit emotional contagion^15^, and display empathy-like behaviors^16^. Indeed, rodent facial expressions and chemosignals, such as olfactory cues, convey internal emotional states and modulate social interactions^17^. Together, these findings highlight the complex socio-emotional repertoire displayed by laboratory animals, which is extremely important to understand the brain substrates controlling social cognition and to tackle potential alterations in these complex social abilities.

Among the brain regions involved in social cognition, amygdala-dependent circuits play a pivotal role in processing socially derived information^18^ and mediating complex behaviors, including observational fear learning^18,19^ and prosocial responses^20,21^. Recent studies have identified specific amygdalar pathways critical for emotion recognition in mice, such as oxytocinergic projections from the hypothalamic paraventricular nucleus to the central amygdala^13^. However, how amygdala circuits contribute to the sex- and age-dependent modulation of emotion recognition remains unclear, despite known influences of sex and aging on amygdalar function^22^.

In this study, we employed an Emotional state Discrimination Task (EDT), a validated paradigm assessing rodents’ ability to discriminate emotionally altered from neutral conspecifics^12–14^. Using TRAP2:Ai14 mice^23^ to selectively capture neurons activated during emotion recognition, combined with genetic, imaging, and chemogenetic approaches, we examined cortico-amygdala circuit engagement. Additionally, we extended our findings to humans using a face recognition task, which demonstrates conserved neural mechanisms across species. Overall, our results reveal sex- and age-dependent cortico-amygdala circuits controlling emotion recognition, highlighting the dynamic nature of socio-affective brain networks and providing insight into circuit-level vulnerabilities underlying socio-cognitive dysfunction across the lifespan.

## Material and Methods

### Animals

Male and female C57BL/6J, Targeted Recombination in Active Populations (TRAP2) and TRAP2:Ai14 mice, aged 8-14 weeks (young mice) and 9-12 months (aged mice) were employed in this study. C57BL/6J mice were obtained directly from Charles River Laboratories (Spain). TRAP2:Ai14 mice were generated by crossing homozygous TRAP2 mice (Fos^tm2.1(icre/ERT2)Luo^/J; The Jackson Laboratory, Bar Harbor, ME, USA; stock #030323) with homozygous *Ai14* tdTomato reporter mice (B6.Cg-Gt(ROSA)26Sor^tm14(CAG-tdTomato)Hze^/J; The Jackson Laboratory; stock #007914). Offspring heterozygous for both transgenes were genotyped as performed in previous studies^23^. All animals were group-housed (2-5 per cage) in a specific pathogen-free facility within a controlled environment (20-24°C and 40-70% of humidity). They had *ad libitum* access to food and water, standard environmental enrichment including nesting material and cardboard houses, and were subjected to a 12-hour light/dark cycle (with lights on at 7:30 pm and off at 7:30 am). Experiments were conducted during the dark phase (between 9 am and 3 pm). Throughout testing, experimenters remained blinded to experimental conditions. All the procedures are adhered to the guidelines of the European Directive on the protection of animals used for scientific purposes (2010/62/EU) and approved by the Animal Ethics Committee of the Parc de Recerca Biomedica de Barcelona (PRBB) and from the Generalitat de Catalunya.

### Drug preparation

4-hydroxytamoxifen (4-OHT; Sigma, Cat# H6278) was prepared as previously described^24^. Briefly, 4-OHT was dissolved at 20 mg/mL in ethanol by shaking at 37°C for 5 min. Before use, 4-OHT was dissolved in Kolliphor oil (Merk, Cat # C5135-500G) by shaking at 37°C for 5 min after which ethanol was evaporated. Finally, Phosphate Buffer (PBS) 0,1M was added to obtain a final 25 mg/kg dose for neuronal activation studies or 50 mg/kg for neuroanatomical tracing experiments. 4-OHT injections were delivered intraperitoneally (i.p) upon completion of the behavioral test.

J60 dihydrochloride ligand (J60; HelloBio, #HB6261) was prepared by dissolving 1 mg of J60 in 100ml of sterile saline solution to reach a concentration of 0,1 mg/kg. The solution was then aliquoted into 15mL Eppendorfs tubes and stored at -20°C until use. The drug was administered intraperitoneally 1 hour prior the behavioral test.

### Surgical procedures and viral vector injections

Animals were anesthetized with an i.p injection of ketamine (75mg/kg) and medetomidine (1mg/kg). They also received a subcutaneous meloxicam injection (5 mg/kg) as analgesic pre-surgery and two days’ post-surgery. After fixing the mouse head in the stereotaxic apparatus and align bregma and lambda coordinates, small holes were drilled in order to infuse viral vectors bilaterally into the basolateral amygdala (BLA) in the following coordinates respect to bregma: AP: -1.5; ML: +/-3,1; DV: -5,2 or into the medial orbitofrontal cortex (MO): AP: +2,6; ML: +/-0,2; DV: -2,3. To study the activity-dependent efferents and afferents from/to the BLA, TRAP2 mice received 250 nL viral injections of the fluorescent anterograde Cre-dependent virus AAV2-hSyn-DIO-mCherry (Addgene # 50459-AAV2, ≥ 4 × 10¹²) or the retrograde Cre-dependent virus rgAAV-hsyn-DIO-EGFP (Addgene #50457-AAVrg, ≥7 × 10¹² vg/mL) bilaterally into the BLA respectively. To investigate the causal involvement of the BLA in emotion recognition C57BL/6J mice received 300 nL of AAV2-CaMKII-hM4D(Gi)-mCherry (Addgene #50477-AAV2, ≥7 × 10¹² vg/mL) or AAV5-CaMKII-mCherry (Addgene #114469-AAV5, ≥7 × 10¹² vg/mL). To study calcium dynamics in the BLA and MO using fiber photometry, C57BL/6J mice were injected with 300 nL of AAV9-CaMKII-GCaMP6m (Tebu-Bi0 #SL101481-AAV9, ≥ 1×10¹³ vg/mL) into the BLA and MO. After virus injection, an optic fiber (core 400 μm, N.A 0.5, RWD, China) was implanted 0.25mm above each viral injection site and was fixed to the skull using dental cement (first layer: superbond, SunMedical; body of the implant). To inhibit the specific projection from MO to BLA during emotion recognition, C57BL/6J mice were co-infused in the BLA with 250 nL of retrograde rgAAV-EF1a-Cre (Addgene #55636-AAVrg, ≥7 × 10¹² vg/mL) and retro rgAAV-CAG-GFP (Addgene #37825-AAVrg, ≥7 × 10¹² vg/mL) at a 4:1 ratio, together with 250 nL of either AAV2-hSyn-DIO-hM4D(Gi)-mCherry (Addgene #44362-AAV2, ≥5 × 10¹² vg/mL) or AAV2-hSyn-DIO-mCherry (Addgene #50459-AAV2, ≥4 × 10¹² vg/mL) in the MO. On the contrary, to inhibit projections from the BLA to the MO, the same viral vectors were employed, but with the injection order inverted. To activate the BLA in aged C57BL/6J mice, 300 nL of the viral vectors AAV5-CaMKII-hM3D(Gq)-mCherry (Addgene #50457-AAV5, ≥ 7 × 10¹² vg/mL) or AAV5-CaMKII-mCherry (Addgene #114469-AAV5, ≥7 × 10¹² vg/mL) were injected into the BLA. After viral vector injections, incisions were sutured, and mice were allowed to recover for 3-4 weeks before the starting of the behavioral protocol.

### Emotional state Discrimination Task

The Emotional state Discrimination Task (EDT) assesses the ability of an observer mouse (experimental subject) to distinguish between a neutral unfamiliar conspecific (neutral demonstrator) and an emotionally altered unfamiliar conspecific (stressed demonstrator), matched for sex, following previously described methodology^12^ with slight variations. The test consisted of three habituation days followed by one test day.

#### Habituation

During habituation, observer mice were placed for 9 min inside a two-chamber maze (6 lux) with an open passage separating the compartments, each containing an empty wire cup. A plastic cylinder was placed on top of the cups to prevent climbing. Demonstrator mice underwent the same habituation procedure inside the cups, without observers present, for 9 min each day.

#### Stress test

On the stress test day, neutral and stressed demonstrators were placed inside the wire cups, and the 6 min test began. On the test day, neutral demonstrators did not receive any manipulation and were left undisturbed, with *ad libitum* water and food access, in their home cage. All neutral demonstrators were group-housed, separately from the cages of stressed demonstrators. Stressed demonstrators were subjected to a mild restraint stress protocol consisting of introducing mice inside a 50 mL falcon tube with holes to allow breathing for 15 min before the test.

#### Neutral test

A neutral test was performed in the experiments using TRAP2:AI14 mice as a control group. This test was performed as previously described for the stress test except that only neutral demonstrators were used.

Demonstrators were test-naive and used a maximum of four times (1 of each demonstrator per 4 observers). The apparatus, cups and separators were cleaned with a combination of ethanol and water and dried with paper towels after changing demonstrators. Moreover, the apparatus and cups were cleaned after each observer when urine was detected. After each day, the wire cups, separators, and cubicles were wiped down with 70% ethanol and allowed to air dry.

### Behavioral quantification

#### Manual scoring

Videos were analyzed offline by experimenter’s blind to group assignments. Time spent sniffing each demonstrator was recorded using BORIS software (v8.22.6). Sniffing was defined as the animal actively directing its nose toward another mouse and making contact or being within close proximity (≤1-2 cm). Raw, % data or a discrimination index (DI) was used for graphical representation of sniffing during the initial 4 minutes of the EDT, which is the period where animals mostly discriminate as shown before^12^. % time sniffing was calculated with the following formula: (TSniff stressed demonstrator/ TSniff stressed demonstrator + TSniff neutral demonstrator) *100 or (TSniff neutral demonstrator/ TSniff neutral demonstrator + TSniff stressed demonstrator) *100. DI was calculated as follows: (TSniff stressed demonstrator – TSniff neutral demonstrator) / (TSniff stressed demonstrator + TSniff neutral demonstrator).

#### Automated scoring

Videos from habituation and test day were preprocessed to homogenize orientation of the behavioral apparatus, frame rate and crop across batches. Subsequently, 14 body parts were tracked automatically using DeepLabCut^25^ and Keypoint-MoSeq^26^ was used to categorize the behavior of mice in an unsupervised manner as recently published by our lab^27^. Regions of interest (ROI) were defined by dividing our square arena in 4 quadrants. Data from the quadrants where the demonstrators were placed were selected for downstream analysis. Custom written python scripts were used to extract metrics from the Deeplabcut output like speed, acceleration, position in the arena, definition of ROI and calculation of DIs using automated metrics like time spent in ROIs (ROI around stressed and neutral animal) or time spent in each Keypoint-MoSeq defined syllable next to the neutral or stressed demonstrator.

### Fiber photometry

#### Behavioral test

We recorded data from the MO and BLA of n=8 male C57BL/6J mice implanted at 8 weeks of age (see surgical procedures above). Experiments were carried out 8 weeks after surgery to allow for expression of GcaMP6m and for healing of the surgery site. Fiber-implanted mice underwent the EDT with a stressed and a neutral demonstrator as described above. They were habituated to plugging and unplugging of the patch cords during the three habituation days.

#### Photometry recording settings

Behavioral and calcium imaging videos were synchronized by using TTL pulses which are built-in in the fiber photometry recording system (RWD). GCaMP fluorescence was recorded at 470nm, and together with it, the isosbestic signal was recorded at 410nm to control for motion artifacts. Behavioral videos were recorded at 20fps and post-hoc down sampled to 10fps. Fiber photometry was recorded at 10fps. From all mice implanted, Mouse 1 lost its MO fiber between surgery and experimental recording and Mice 2 and 3 did not show expression of GCaMP6m under the MO fiber tip and were hence discarded from downstream analysis.

#### Data preprocessing

A custom-written Python script was used to preprocess the data and analyze the behavioral and fluorescence time series. For preprocessing, (1) the first minute of the recording was discarded to avoid the effect of the initial photobleaching that is characteristic in fiber photometry. (2) The 470nM signal was fitted to the isosbestic 410nM using a linear fit and for each time point (3) dF/F0 was calculated as (F470nm – F410nm(fitted))/ F410nm(fitted) and multiplied by 100 to express it as a percentage of change.

### Histology

TRAP2 mice were euthanized four weeks after 4-OHT injections to visualize activity-dependent tracers. TRAP2:Ai14 mice were euthanized two weeks after the behavioral experiment, while C57BL/6J mice were euthanized one week after. In all cases, euthanasia was performed using ketamine (100 mg/kg) and xylazine (20 mg/kg). Transcardial perfusion was then performed using ice-cold 4% paraformaldehyde (PFA) at pH 7.3. The brains were extracted and post-fixed in 4% PFA at 4 °C for 24 hours before being transferred to a 30% sucrose solution for cryoprotection. Once equilibrated, the brains were sectioned at 30 μm using a Leica CM3050 S cryostat. Free-floating brain sections were collected and stored in an antifreeze solution. For histological analysis of reporter genes, brain sections were mounted on glass slides and coverslipped with Fluoromount-G™ mounting media containing DAPI (Invitrogen, Cat #00-4959-52) to visualize tdTomato-expressing cells or viral vector expression. For fiber photometry experiments, brain sections around the target brain areas where viral vectors were infused and fibers implanted were collected for verification of the fiber location and the expression of GCaMP following the above-described process.

To perform immunohistochemistry analysis, brains sections were first washed three times in PB 0.1M followed by an additional wash in PB 0.1M containing 0.2% Triton X-100. The sections were then incubated in a blocking buffer consisting of PB 0.1M with 0.2% Triton X-100 and 5% donkey serum for 60 min. Sections were then incubated with Anti-c-Fos primary antibody (rabbit, 1:1000, Synaptic System, #226 008) overnight at 4 °C. After washing, sections were incubated with a secondary antibody (donkey anti-rabbit, 1:1000, AlexaFluor 488-conjugated # 711-545-152,) for 2 h at room temperature. After extensive washing, the sections were mounted on glass slides using Fluoromount-G™ mounting medium with DAPI.

### Epifluorescence microscopy

Fluorescent images were acquired with a Nikon ECLIPSE Ni-E motorized microscope, using a 10x/0.40NA objective. The Nis ELEMENTS software was used to perform image acquisitions. Image expression of tdTomato, mCherry, GFP, c-Fos, GCaMP and DAPI were examined in tissue containing various anterior-posterior coronal levels of the studied areas.

### Image data analysis

ImageJ software 1.54i (National Institutes of Health, Bethesda, Maryland, USA) was used to quantify tdTomato cells in the sections under fluorescent microscopy. Cells were manually considered to be tdTomato, when showing appropriate size (diameter ranging approximately 20 µm) and distinct from the background at 10× magnification. A custom macro was developed to detect c-Fos+ neurons. The different analysis involved around 3-5 photos per mouse from each area, and the average of these images are represented in the graphics. The bregma coordinates of each area are the following: VO (2.80-2.46), MO (2.80-2.46), Cg (1.78-1.54), PrL (1.78-1.54), NAc (1.78-1.54), POA (0.14-0.02), AIP (0.14-(-0.22)), pBNST ((-0.22)-(-0.34)), PVH ((-0.58)-(-1.22)), LHb ((-0.58)-(-1.22)), PVT ((-0.82)-(-0.34)) and BLA/CeA ((-1.22)-(-1.58)).

### Human data

#### Participants

The dataset is part of a population-based neuroimaging-genetics study: the IMAGEN cohort^28^. For this study, we analyzed neuroimaging data from 1417 participants at 19 years old. After quality control procedure, 1263 participants (age 19.08 ± 0.76 years old, 52.7% female) were included in the analysis. Participants were recruited from eight research centers in the United Kingdom, German, France and Ireland. Ethics approval was obtained at each site by local research ethics committee. Written consent was obtained from each participant.

#### Emotional faces task (EFT) fMRI paradigm

The study used an fMRI paradigm based on previous studies^29^ to investigate social-emotional processing. Participants viewed 18-second blocks of either face stimuli (angry, happy, or neutral faces) or non-face stimuli (control). The face stimuli consisted of black-and-white video clips (2-5 seconds each) showing moving faces from three male and three female actors. The control stimuli were black-and-white concentric circles expanding or contracting, designed to match the visual contrast and motion of the face clips. Each type of face stimulus was presented four times, interspersed with twelve blocks of the control stimuli. The total scanning session for the task was approximately 7 minutes long.

#### MRI acquisition and preprocessing

Neuroimaging data were collected from eight different study centers using 3T scanners. The scanning parameters were harmonized across all sites. Visual stimulus was presented using the standardized hardware (Nordic Neurolabs, Bergen Norway). For each participant, two types of scans were performed: a high-resolution T1-weighted MRI (T1) scan using the Magnetization Prepared Rapid Acquisition Gradient Echo (MPRAGE) sequence with the following parameters: repetition time (TR) = 2,300 ms; echo time (TE) = 2.8 ms; flip angle (FA) = 8°; isotropic voxel size 1.1 mm; 256 × 256 × 160 matrix; sagittal slice plane. Functional images were acquired using gradient-echo, echo-planar imaging (EPI) sequence with the parameters: TR = 2,200 ms, TE = 30 ms, FA = 75°, voxel size=3.4 × 3.4 × 2.4 mm, slice ga*p =* 1mm and 64 × 64 × 40 matrix.

Task-based fMRI scans were pre-processed using the Statistical Parametric Mapping (SPM12, http://www.fil.ion.ucl.ac.uk/spm/). The pre-processing steps included: non-brain tissue removal; slice-timing correction to account for the different acquisition times of each brain slice; head movement realignment to correct for participant motion during the scan and non-linearly warping of images to the standard MNI space using a customized EPI template. Spatial smoothing of normalized images using a 5mm FWHM Gaussian kernel. Pre-processed data were resliced to 3mm isotropic voxels. To ensure signal stability and scanner equilibrium, the first five volumes were discarded. Region of interest (ROI) was defined based on automated anatomical labelling atlas 3 (AAL3)^30^.

##### ROI-Based Activation Analysis

We analyzed human EFT fMRI data from the IMAGEN cohort. Beta coefficients representing activation amplitude were extracted from three bilateral ROIs: the amygdala, ventromedial prefrontal cortex (vmPFC), which is often considered a putative homolog to the MO cortical region of mice^31^, and other subregions of the human orbitofrontal cortex (OFC) (Fig. 1A). For each ROI, outliers exceeding median ± 3 interquartile ranges were excluded. We then calculated the angry vs. neutral contrast by subtracting the beta coefficients of the neutral condition from those of the angry condition. To assess sex differences in this angry vs. neutral contrast, we used linear regression models including imaging site and handedness as covariates.

**Figure 1.**
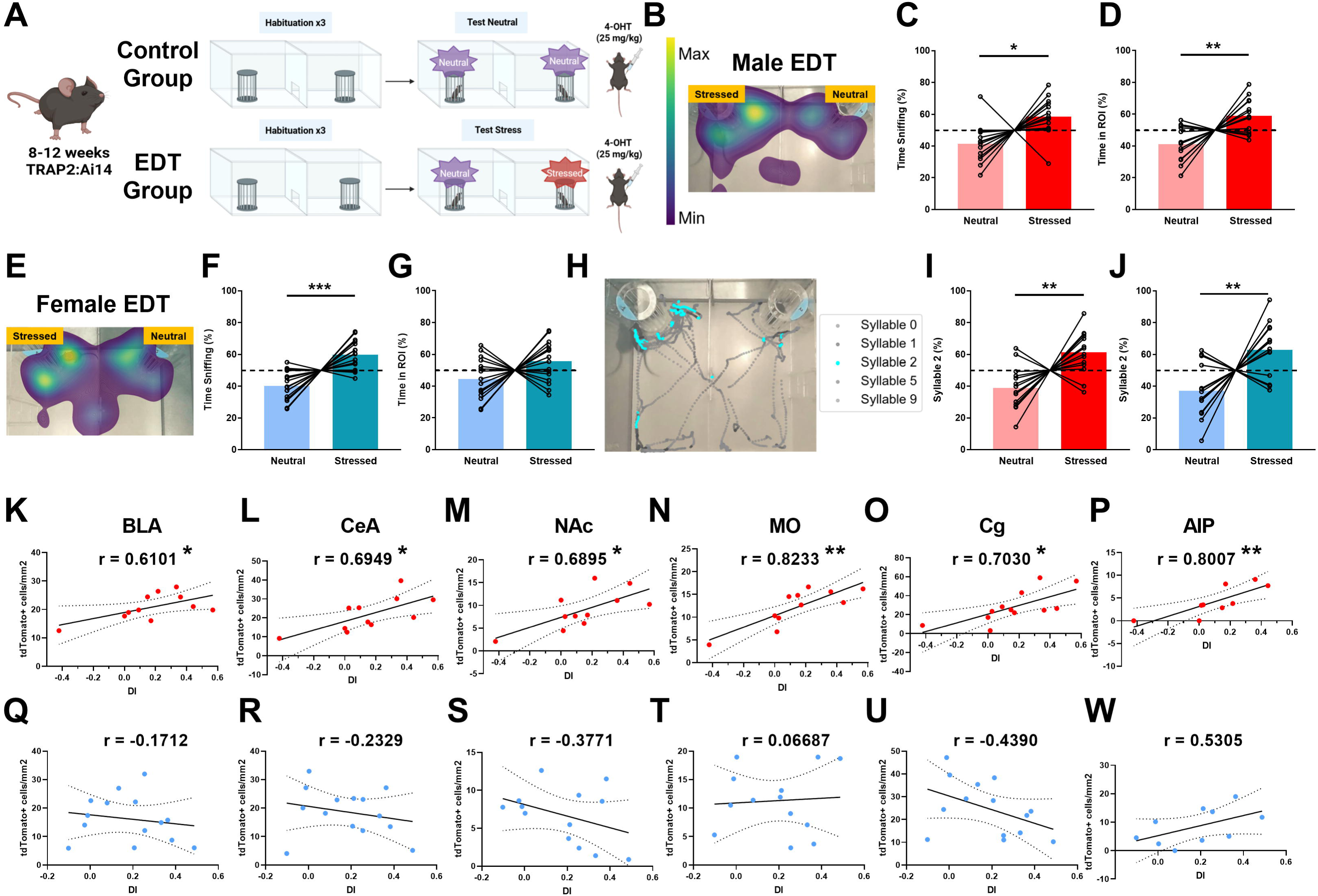
Sex-dependent engagement of brain circuits during emotion recognition in young mice. (A) Schematic representation of the experimental design used in the negative Emotional state Discrimination Task (nEDT) in young TRAP2:Ai14 mice (8-12 weeks old), showing control and EDT groups. (B) Representative heatmap showing exploration patterns of male TRAP2:Ai14 mice during the nEDT. (C) Percentage of time spent sniffing the neutral versus stressed demonstrator during the task in young males. Dashed indicate the 50% chance level used as a reference in one-sample *t*-test analyses (D) Percentage of time spent in a region of interest (ROI) adjacent to the neutral or stressed demonstrator’s compartment during the nEDT in young males. (E) Representative heatmap of female TRAP2:Ai14 mice during the nEDT. (F) Percentage of time spent sniffing the neutral versus stressed demonstrator during the task in female mice. (G) Percentage of time spent in the ROI adjacent to the neutral or stressed demonstrator’s compartment during the task in female mice. (H) Behavioral representation showing enrichment of behavioral syllable 2 (in blue) identified using the unsupervised Keypoint-MoSeq algorithm. (I-J) Percentage of time spent expressing behavioral syllable 2 only when interacting in the neutral or stressed demonstrator ROI during the task in males (I) and females (J). (K-P) Correlation analyses between tdTomato+ cells per mm^2^ and DI (time spent sniffing neutral vs. stressed demonstrator) in young male TRAP2:Ai14 mice for the BLA (K), CeA (L), NAc (M), MO (N), Cg (O), and AIP (P). (Q-W) Equivalent correlation analyses in young female TRAP2:Ai14 mice for the BLA (Q), CeA (R), NAc (S), MO (T), Cg (U), and AIP (W). BLA: Basolateral Amygdala; CeA: Central Amygdala; NAc: Nucleus Accumbens; Cg: Cingulate cortex; MO: Medial Orbitofrontal cortex; AIP: Agranular Posterior Insular cortex. Data are represented as before-after for the percentage of time graphs. In the correlations, lines represent linear regression fits; shaded areas show 95% confidence intervals. ***P < 0.001; **P < 0.01; *P < 0.05. For statistical details, see SI Appendix, Table S1.

##### Functional Connectivity Analysis

Task-based connectivity was examined using beta-series correlation analysis^32^ with the bilateral amygdala as the seed region. For each participant, we computed Pearson correlations between the amygdala beta coefficient and whole-brain voxel-wise beta coefficients separately for angry and neutral conditions.

### Statistical methods

All statistical analyses were performed with GraphPad Prism 8.0 software and with custom-written Python scripts. Data are presented as mean ± standard error mean (SEM). Normality was assessed with a Shapiro-Wilk test. For data following a normal distribution, between-group comparisons were performed by paired t-test, one sample t-test or two-way ANOVA analysis in behavioral task and one way ANOVA or unpaired t-test for image quantification. When normality was not met, nonparametric Kruskal Wallis or Mann-Whitney U test were used (Supplementary Material). Outliers, defined as values exceeding two SD above or below the mean of the experimental condition, were identified and removed. A p-value of <0.05 was considered as significant. Regarding the photometry analysis, all analysis and visualizations were carried out by custom-written python scripts. Average Z-score comparisons between time spent sniffing the stressed, the neutral demonstrators or other behaviors (Figure 3, F) were carried out by selecting the average Z-score of all the frames where each behavior happened in every mouse and comparing the averages with a paired t-test independently in BLA and OFC. Z-score was calculated on the entire recording by subtracting the median of the ΔF/F₀ to the ΔF/F₀ and dividing by the mean absolute deviation of the ΔF/F₀. Analysis focused on the onset and offset of sniffing of the demonstrators (Figure 3, G-L) was carried following these steps (1) trials were defined as sniffing bouts that lasted for >1s. (2) fluorescence was selected in -/+2s windows around trial onset and offset. (3) each trial was normalized by computing the z-score with the median of the first second of the trial (baseline), subtracting it to it and dividing by the mean absolute deviation of the baseline. Statical significance was assessed region by region and separating trials were mice explored either the neutral or stressed conspecific by ranking the maximum value of the average trial across mice in the distribution of maximum values of 10.000 randomly generated trials. The temporal and trial structure of the real data was preserved in the bootstrapped data. Statistical significance was determined by assessing which Z-score values are greater than the 95% confidence interval of the bootstrapped distribution. Finally, regarding the human analysis, we used paired t-tests to examine the difference between angry and neutral conditions within each sex in the ROI-Based activation analysis. We corrected for multiple comparisons across the ROIs using the False Discovery Rate (FDR), with a significance threshold of *q* < 0.05. On the other hand, one-sample t-tests were used to identify significant connectivity difference between angry and neutral conditions. Voxel-wise significance threshold was set at *p* < 0.001 for the group-level results.

## Results

### Sex-dependent engagement of brain circuits during emotion recognition in young mice

To investigate the neural mechanisms underlying emotion recognition, we used a validated negative Emotional state Discrimination Task (nEDT) in mice, where an “observer” discriminates between an unfamiliar neutral “demonstrator” mice and an unfamiliar stressed demonstrator (Fig. 1A)^12–14^. The negative state of the stressed demonstrator was induced via mild restraint prior to the task^12–14^. To map brain regions activated during this task, we employed TRAP2:Ai14 mice^23^, which allow for activity-dependent tagging of neurons. Two experimental groups were used (Fig. 1A): (i) the EDT group, exposed to a stressed and a neutral demonstrator; and (ii) the Control group, exposed to two neutral demonstrators. No bias was observed for the neutral demonstrator placed on the left or the right side of the maze (Suppl. Fig. 1A-B), indicating no spatial bias in the EDT arena used. Mice received 4-hydroxytamoxifen (4-OHT) immediately after the task to label activated neurons, and brains were collected two weeks later. As previously shown in C57BL/6J mice^12–14^, male TRAP2:Ai14 mice showed increased sniffing (Fig. 1B-C, Suppl. Fig. 1C) and time spent near the stressed demonstrator (Fig. 1D). Females displayed higher locomotion (Suppl. Fig. 1D-E),but similarly showed more sniffing (Fig. 1E-F, Suppl. Fig. 1F) and proximity to the stressed mouse (Fig. 1G), confirming emotion discrimination in both sexes. To gain deeper behavioral insights, we used DeepLabCut^25^ for pose estimation and Keypoint-MoSeq^26^ for unsupervised behavioral classification. This analysis revealed that a specific behavioral stereotyped action (“syllables 2”; Fig. 1H), correlated with manually scored sniffing (Suppl. Fig. 1G-H). Indeed, syllable 2 was more frequent in the stressed compartment for both sexes (Fig. 1I-J), supporting its role as a high-resolution, unbiased marker of social investigation that could be used to dissect social investigation dynamics in EDT or similar social behavioral tasks.

To investigate the brain regions engaged during the nEDT, we analyzed the brains from TRAP2:Ai14 mice after the behavioral test. This approach allowed us to examine whether regional activity levels (*i.e*., tdTomato expression) were correlated with emotion recognition performance (*i.e*., Discrimination index; DI). In male mice, tdTomato expression in the BLA, Central Amygdala (CeA), Nucleus Accumbens (NAc), Medial Orbitofrontal cortex (MO), Cingulate cortex (Cg), and Anterior Insula (AIP) positively correlated with nEDT performance (Fig. 1K-P, Suppl. Fig. 1I-J). Strikingly, no such correlations were found in female mice (Fig. 1Q-W, Suppl. Fig. 1I and 1K) or in the control group (Suppl. Fig. 1L-M). Together, these results suggest that despite presenting similar behavioral performance, male and female mice may be recruiting sex-specific circuits to support emotion recognition.

### Dynamic BLA and MO engagement during negative emotion recognition

Given the privileged position of the amygdala at the interface between cortical and subcortical networks, and its well-established role in emotional processing, we first mapped BLA-related circuits engaged during emotion recognition. To this end, we injected Cre-dependent retrograde and anterograde AAVs expressing GFP and mCherry into the BLA of TRAP2 mice to dissect activity-dependent projections recruited during the nEDT (Suppl. Fig. 2A). Post-task anatomical analysis revealed activated bidirectional cortico-amygdalar projections during the nEDT, including the MO, Prelimbic cortex (PrL), and AIP (Suppl. Fig. 2B-C), supporting their potential involvement in emotion recognition^12,18,20,33^.

Since the contributions of the PrL and insular cortex to emotion recognition have been previously demonstrated in rodents^12,20,34,35^, we focused our attention on the MO in order to explore the potential involvement of specific orbitofrontal cortex (OFC) subregions and how they interact with the BLA during emotion recognition. Therefore, to assess real-time dynamics in BLA and MO during the nEDT, we performed fiber photometry in male C57BL/6J mice infused with AAV-CaMKII-GcaMP6m into these brain regions (Fig. 2A-D, Suppl. Fig. 2D-F). Both regions showed increased activity during active social exploration vs. baseline (Fig. 2E-F), and distinct activation peaks when mice initiated (Fig. 2G-I) or ended (Fig. 2J-L) sniffing towards the conspecifics. These findings confirm dynamic BLA-MO engagement during social investigation in a nEDT, consistent with our prior anatomical data obtained with the TRAP2 mice, and identify a temporal engagement of both brain regions when exploring conspecifics with different internal states.

**Figure 2.**
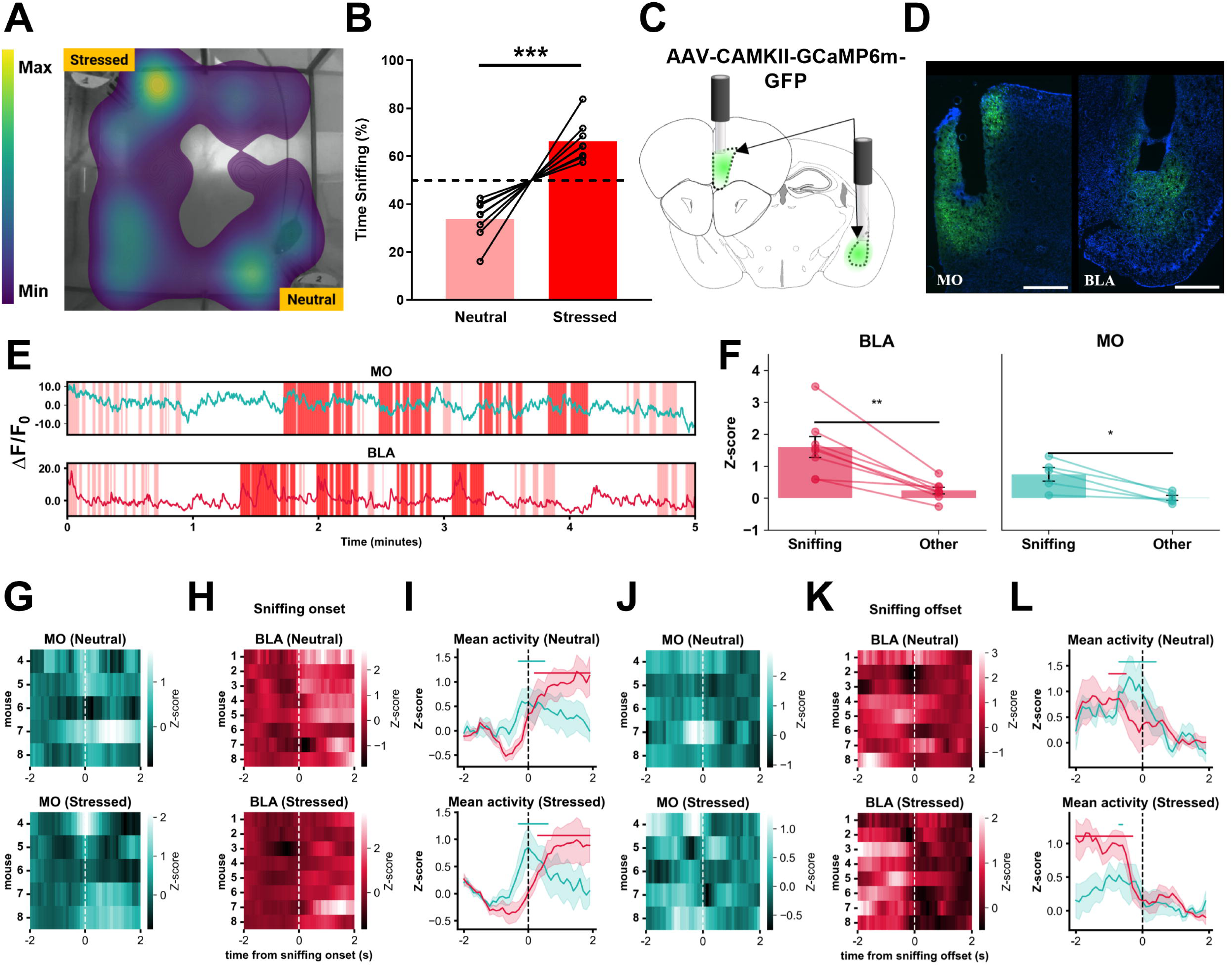
Dynamic BLA and MO engagement during negative emotion recognition. (A) Exemplary heatmap showing time density in the arena. (B) Fiber-implanted mice spent more time sniffing a stressed demonstrator than a neutral one. (C) Schematic showing surgery set up. (D) Representative histologies showing fiber placement and GCaMP6m injections in MO (left) and BLA (right). (E) Representative ΔF/F₀ over time for MO (top) and BLA (bottom). Colored patches indicate sniffing of the neutral (pink) or stressed (red) conspecifics. (F) Average Z-score for all mice during sniffing the demonstrators and other behaviors for BLA (left) and MO (right). Statistical significance tested paired t-test. (G) Average ΔF/F₀ in MO around sniffing onset for all implanted mice for sniffing the neutral (top) or the stressed (bottom) conspecific. (H) Average ΔF/F₀ in BLA around sniffing onset for all implanted mice for sniffing the neutral (top) or the stressed (bottom) conspecific . (I) Average MO and BLA traces for all mice. Significance tested by assessing time windows in which Z-score was highet than the 95% confidence intervalof adistribution of 10000 randomly sampled average traces (bootstrapping). Significant time windows are indicated with horizontal bars colored accoridng to the brain region. (J-L) Equivalent graphs as G,H and I for offset of sniffing. Fluorescent images represent 10x of GCaMP-GFP signal. (Scale bar, 500 μm). ***P < 0.001; **P < 0.01; *P < 0.05. For statistical details, see SI Appendix, Table S1.

### Reciprocal BLA-MO projections mediate emotion recognition in young mice

To test causality, we used chemogenetics to inhibit BLA principal neurons in male and female C57BL/6J mice during the nEDT (Fig. 3A-B). In male mice, BLA inhibition abolished emotion recognition (Fig. 3C, Suppl. Fig. 3A) without affecting locomotion or sniffing behavior (Suppl. Fig. 3B-D). In contrast, female mice retained emotion discrimination (Fig. 3D, Suppl. Fig. 4E-H), indicating a sex-dependent involvement of the BLA during emotion recognition. Next, we employed an intersectional viral approach to selectively inhibit bidirectional projections between the MO and BLA (Fig. 3E-H). Inhibiting either MO-BLA or BLA-MO projections disrupted emotion recognition in male mice (Fig. 3I, Suppl. Fig. 3I) but had no effect in female mice (Fig. 3J, Suppl. Fig. 3M), further supporting a sex-specific brain circuit involved in negative emotion recognition. Locomotor behavior was mostly unaffected (Suppl. Fig. 3J-P), although BLA-MO inhibition increased total sniffing in males (Suppl. Fig. 3J, 3N). These results reveal that bidirectional BLA-MO communication is necessary for emotion recognition in male, but not, female mice.

**Figure 3.**
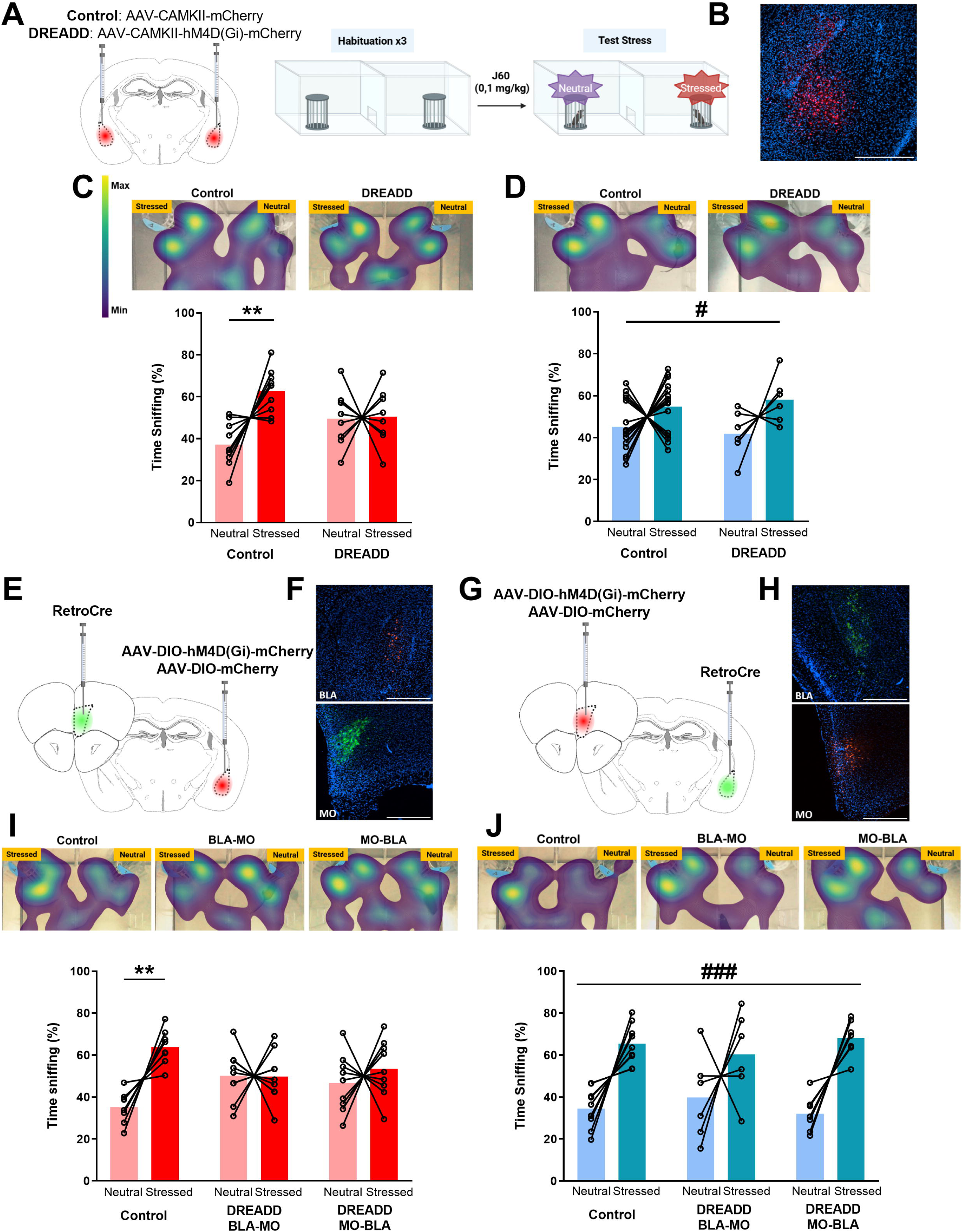
Chemogenetic inhibition of the BLA and reciprocal BLA-MO projections during emotion recognition. (A) Schematic representation of bilateral infusions of control virus (AAV-CAMKII-mCherry) or inhibitory DREADD (AAV-CAMKII-hM4D(Gi)-mCherry) into the BLA of young male and female C57BL/6J mice, followed by intraperitoneal injections of J60 (0.1 mg/kg) 1 h prior to the nEDT. (B) Representative image of BLA showing inhibitory DREADD-mCherry expression. (C) Top: Representative heatmaps illustrating exploration patterns of control (left) and DREADD (right) male mice during the nEDT. Bottom: Percentage of time spent sniffing neutral versus stressed demonstrators in control and DREADD male groups. (D) Top: Representative heatmaps showing exploration patterns of control (left) and DREADD (right) female mice during the nEDT. Bottom: Percentage of time spent sniffing neutral versus stressed demonstrators in control and DREADD female groups. (E-F) Schematic (E) of intersectional viral strategy to selectively inhibit BLA-MO projections using retrograde Cre delivery to MO combined with Cre-dependent inhibitory DREADD in the BLA. (F) Representative images of retrograde viral expression (green) in MO and inhibitory DREADD expression (red) in the BLA. (G-H) Schematic (G) of viral strategy to inhibit MO-BLA projections using retrograde Cre delivery to the BLA and Cre-dependent inhibitory DREADD expression in MO. (H) Representative images showing retrograde viral expression (green) in the BLA and inhibitory DREADD expression (red) in MO. (I) Top: Representative heatmaps of exploration patterns in male mice across Control (left), BLA-MO inhibition (middle), and MO-BLA inhibition (right) groups. Bottom: Percentage time sniffing neutral versus stressed demonstrators in male experimental groups. (J) Top: Representative heatmaps of exploration patterns in female mice across Control (left), BLA-MO inhibition (middle), and MO-BLA inhibition (right) groups. Bottom: Percentage time sniffing neutral versus stressed demonstrators in female experimental groups. Fluorescent images represent 10x of mCherry or GFP signal expression. (Scale bar, 500 μm). Data are represented as before-after for the percentage of time sniffing. In the repeated measure two-way ANOVA statistical analysis. * denote significant interaction between the two factors while # denote significant effect of demonstrator factor (neutral versus stressed). ***/### P < 0.001; **/## P < 0.01; */# P < 0.05. For statistical details, see SI Appendix, Table S1.

### Amygdala and orbitofrontal cortex activity during emotional face processing in human subjects

To evaluate conservation of these circuits in humans, we used fMRI during an emotional faces task (EFT) (Fig. 4A)^29^. In males, angry vs. neutral faces evoked greater activation in the amygdala and posterior OFC, while the ventromedial prefrontal cortex (vmPFC) showed reduced deactivation (Fig. 4B). In females, only vmPFC deactivation remained significant^28^; neither amygdala nor OFC activation survived correction. Direct sex comparison showed stronger amygdala activation in males, with a nominal (but non-significant after correction) sex effect in the OFC. Interestingly, functional connectivity analysis revealed that males exhibited increased vmPFC-amygdala connectivity, while females showed enhanced OFC-amygdala coupling (Fig. 4C). Despite a full anatomical correspondence between mouse and human orbitofrontal subregions cannot be attributed, these findings suggest an evolutionary conserved sex-specific recruitment of cortico-amygdala circuits during emotion recognition across different species.

**Figure 4.**
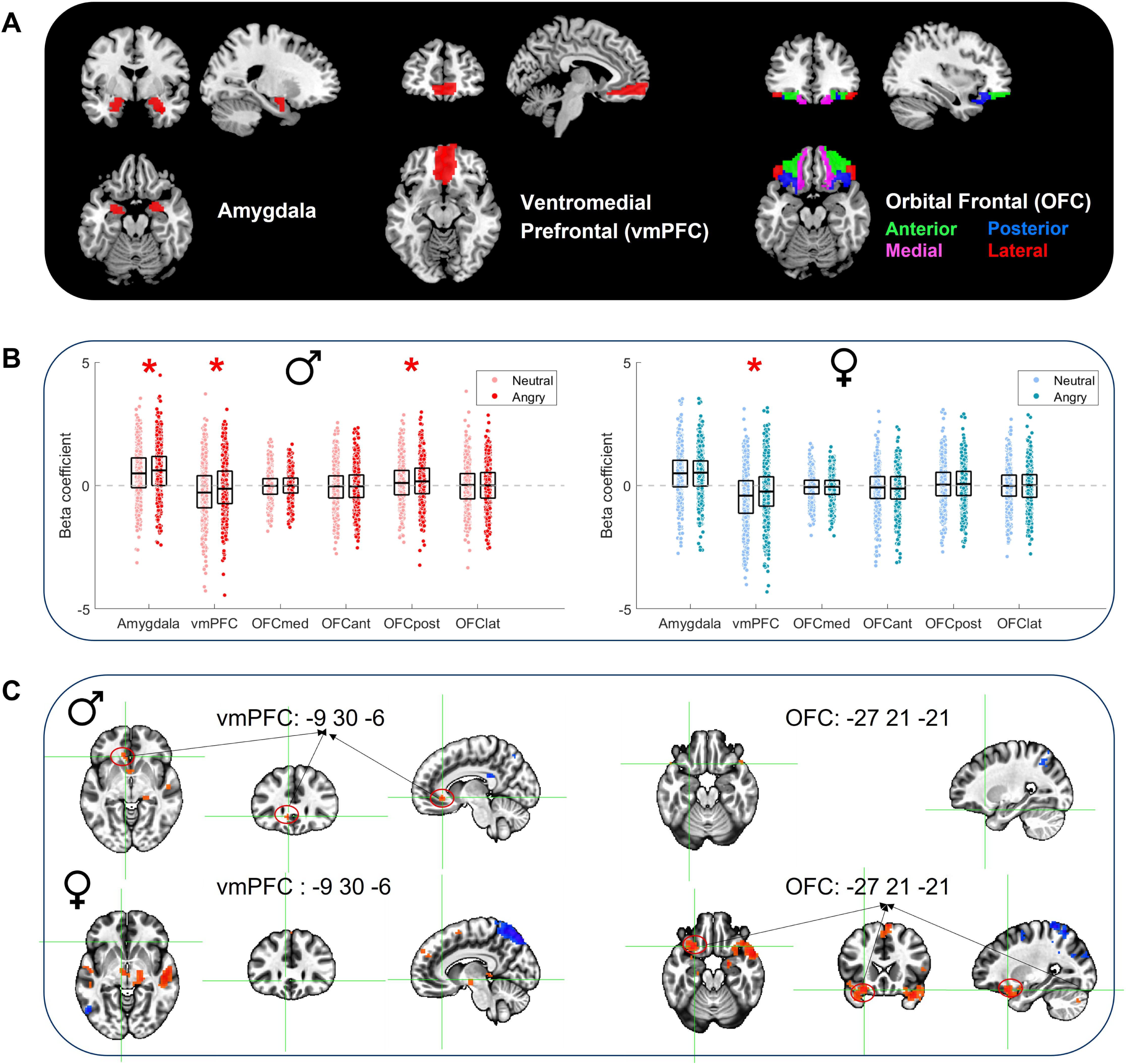
Brain activation and connectivity during human emotional face processing. (A) Anatomical locations of the ROIs examined in this study. Six bilateral ROIs are displayed on template brain images: amygdala, ventromedial prefrontal cortex (vmPFC), and four subdivisions of the orbitofrontal cortex (OFC) -anterior (green), posterior (bule), medial (pink), and lateral (red). ROIs are shown in coronal, sagittal, and axial views. (B) Distribution of beta coefficients for angry and neutral faces conditions across all six ROIs, separated by sex. Left panel shows males (♂), and the right panel shows females (♀). Asterisks (*) denote significant differences between angry and neutral faces conditions (p < 0.05, FDR-corrected). (C) T-statistic maps showing significant connectivity differences between angry and neutral faces conditions, using the bilateral amygdala as the seed region. Red clusters indicate stronger amygdala connectivity during angry relative to neutral faces, whereas blue clusters indicate the opposite. Males (top) show selective vmPFC-amygdala connectivity, while females (bottom) demonstrate OFC-amygdala connectivity. Cross-hairs indicate peak coordinates for each sex-specific cluster (p < 0.001). For statistical details, see SI Appendix, Table S1.

### Age-dependent involvement of amygdala in emotion recognition

We next assessed whether age-related deficits in emotion recognition are linked to altered amygdalar function^36,37^. To test this, aged (9-12 months) TRAP2:Ai14 mice underwent the nEDT (Fig. 5A). Notably, aged male mice failed to show preferential sniffing of stressed demonstrators, suggesting impaired emotion discrimination (Fig. 5B, Suppl. Fig. 4A), while aged female mice conserved this ability (Fig. 5C, Suppl. Fig. 4B). No side or locomotor biases were observed in these experiments (Suppl. Fig. 4C-F). Remarkably, no task-induced activation was observed in BLA or MO, and activity did not correlate with performance in either sex (Fig. 5D-K, Suppl. Fig. 4G-H), in contrast to the patterns observed in young males (Fig. 1). These data suggest that age-related deficits in males are associated with diminished recruitment of key emotional processing regions. To test whether restoring BLA activity could rescue the emotion recognition deficit observed in aged male mice, we expressed excitatory DREADDs in the BLA of aged mice (Fig. 5L, Suppl. Fig. 4I-K). Interestingly, targeted BLA activation rescued emotion recognition in aged males (Fig. 5M, Suppl. Fig. 4L) without affecting general behavior (Suppl. Fig. 4M-O). No effects were observed in aged females (Fig. 5N, Suppl. Fig. 4P-S), confirming the causal role of BLA in male-specific emotion discrimination and its decline with aging.

**Figure 5.**
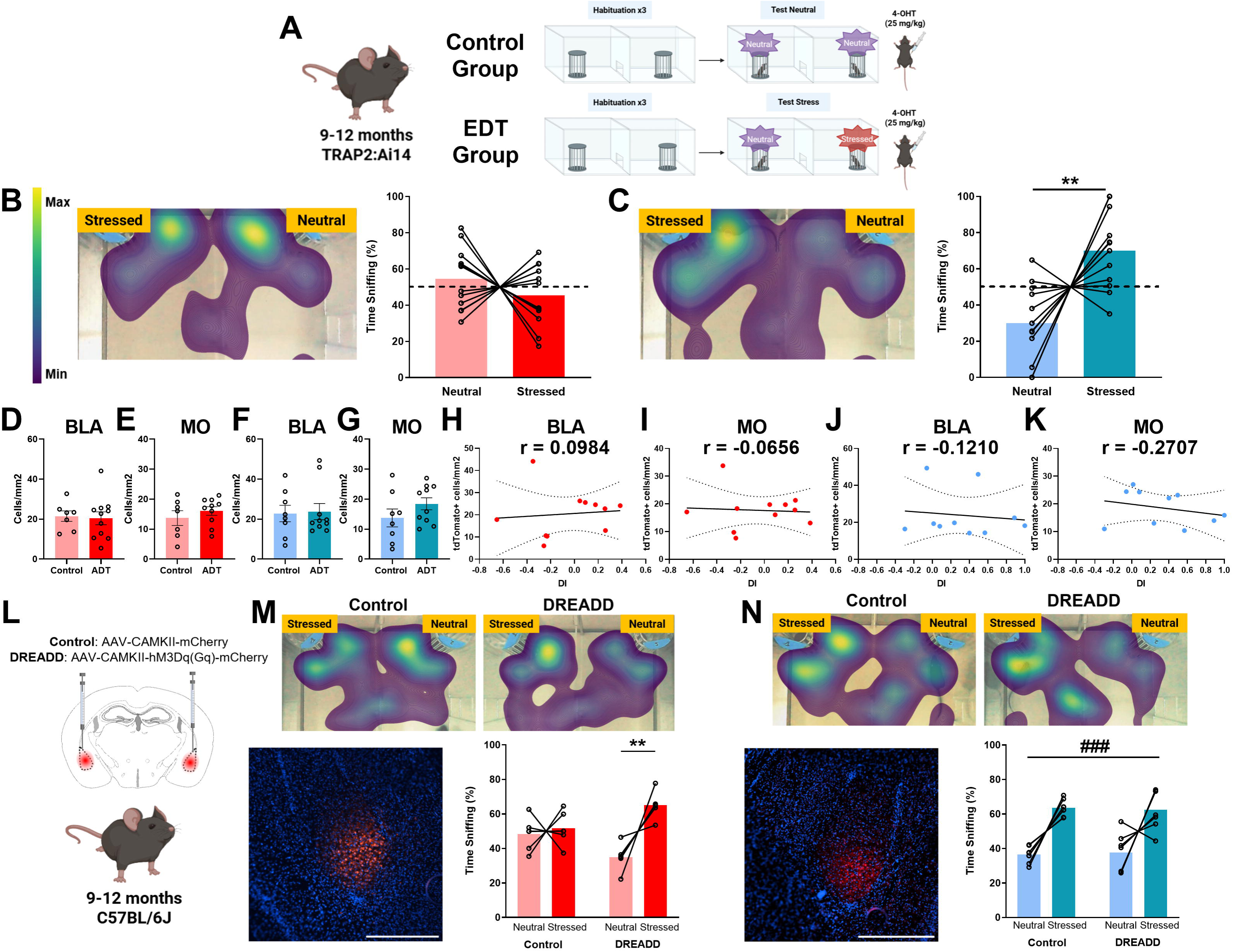
Age-dependent involvement of amygdala in emotion recognition. (A) Schematic of the experimental design for the nEDT in aged TRAP2:Ai14 mice (9-12 months old), showing control and EDT groups. (B) Left: Representative heatmap illustrating exploration patterns of aged male TRAP2:Ai14 mice during the nEDT. Right: Percentage of time spent sniffing neutral versus stressed demonstrators in aged males. (C) Left: Representative heatmap of exploration patterns of aged female TRAP2:Ai14 mice during the nEDT. Right: Percentage of time spent sniffing neutral versus stressed demonstrators in aged females. (D-E) Quantification of tdTomato+ cells per mm² in the BLA (D) and MO (E) between Control and EDT aged male TRAP2:Ai14 groups. (F-G) Quantification of tdTomato+ cells per mm² in the BLA (F) and MO (G) between Control and EDT aged female TRAP2:Ai14 groups. (H-I) Correlations between tdTomato+ cell density and time sniffing DI in aged male TRAP2:Ai14 mice for the BLA (H) and MO (I). (J-K) Correlation between tdTomato+ cell density and time sniffing DI in aged female TRAP2:Ai14 mice for the BLA (J) and MO (K). (L) Schematic of bilateral viral infusions of control (AAV-CaMKII-mCherry) or excitatory DREADD (AAV-CaMKII-hM3Dq-mCherry) into the BLA of aged male and female C57BL/6J mice, followed by intraperitoneal J60 (0.1 mg/kg) injection 1 h before the nEDT. (M) Top: Representative heatmaps of exploration patterns in control (left) and DREADD (right) aged male mice during the nEDT. Bottom left: Representative image of BLA showing excitatory DREADD-mCherry expression. Bottom right: Percentage time spent sniffing neutral versus stressed demonstrators in control and DREADD aged male groups. (N) Top: Representative heatmaps of exploration patterns in control (left) and DREADD (right) aged female mice during the nEDT. Bottom left: Representative image of BLA showing excitatory DREADD-mCherry expression. Bottom right: Percentage time spent sniffing neutral versus stressed demonstrators in control and DREADD aged female groups. Fluorescent images represent 10x of mCherry signal expression. (Scale bar, 500 μm). Data are represented as before-after for the percentage of time sniffing or as mean ± SEM for the quantification of tdTomato+ cells. Dashed lines in the %time sniffing graph indicate the 50% chance level used as a reference in one-sample *t*-test analyses. In the correlations analysis lines represent linear regression fits; shaded areas show 95% confidence intervals. In the repeated measure two-way ANOVA statistical analysis * denote significant interaction between the two factors while # denote significant effect of demonstrator factor (neutral versus stressed). ***/### P < 0.001; **/## P < 0.01; */# P < 0.05. For statistical details, see SI Appendix, Table S1.

## Discussion

Our study uncovers a previously unrecognized, evolutionarily conserved, and sex-dependent brain circuit, comprising bidirectional cortico-amygdala projections, that regulates emotion recognition in both mice and humans. We further demonstrate that the ability to recognize emotionally altered conspecifics declines with age in a sex-dependent manner and that activating the BLA during emotion recognition rescues this deficit. These findings establish a causal role for BLA-centered circuits in supporting emotion recognition and highlight the BLA as a potential therapeutic target in disorders characterized by impaired emotion recognition.

Consistent with previous studies^12–14^, both male and female mice could discriminate between stressed and neutral conspecifics. However, other studies using different behavioral approaches have reported enhanced emotion discrimination in female rodents^10,38^. This variability may stem from the emotional salience of the stimuli as sex differences are more consistently found when strong aversive triggers (e.g., pain^39,40^, footshocks^38^) are used, but not in paradigms using milder stressors such as acute restraint^12–14^. Such enhanced sensitivity in females has been proposed to reflect adaptive evolutionary pressures related to offspring protection^41^.

Our data reveal a sex-and performance-dependent engagement of specific brain regions. In young male mice, emotion recognition performance is correlated with the engagement of a distributed network of cortical and subcortical structures, a network differentially engaged in female mice. These sex differences may be explained by (i) the recruitment of distinct circuits in females while preforming the EDT, (ii) sex-specific differences in 4-OHT metabolism^42^ or baseline expression of immediate-early genes^43^, or (iii) hormonal fluctuations influencing circuit activation^44^. Supporting the latter, studies in women show enhanced amygdala and OFC activation and improved emotion recognition during specific hormonal phases, particularly in response to negative stimuli^45,46^.

Among the regions engaged, the amygdala, specifically the BLA and CeA, emerged as a central hub. Its activation in male mice, but not female mice, may be linked to the use of mild restraint stress to induce emotional changes in demonstrators^47^, consistent with previous human findings^48^. Beyond the amygdala, male emotion recognition was associated with activation in PFC subregions (MO, VO, PrL, Cg), the AIP, and the NAc. These regions have been widely implicated in social cognition and emotion processing^12,49,50^. Notably, activation in the AIP positively correlated with behavioral performance, aligning with studies linking insular damage to deficits in emotion recognition^51^. Although dissecting basal ganglia contributions in humans is limited by imaging resolution, our rodent data suggest a sex-specific role for the NAc in processing socially relevant emotional cues, expanding current views on basal ganglia involvement in socio-cognitive behavior^52^. Overall, our data exemplify an incredible complex puzzle underlying negative emotion recognition in mice that aligns with several key regions observed in human subjects.

Our fiber photometry recordings confirmed that both BLA and MO are dynamically engaged during emotion recognition, with MO activation peaking during the initiation of social exploration and BLA activation increasing once the interaction is established. These temporal dynamics suggest functional coordination during emotion processing, regardless of the demonstrator’s emotional state. Higher-resolution tools may help to further dissect these subtleties, including potential cell-type specific contributions. Indeed, using chemogenetics, we demonstrated that bidirectional BLA-MO projections are required for emotion recognition in male mice. BLA-MO signaling may convey emotional valence, helping assign significance to observed cues^53^, while MO-BLA projections may regulate the salience of these cues to shape appropriate behavioral responses^54^. Disrupting either projection impaired emotion recognition without affecting general exploration, confirming the specificity of this circuit. Our findings align with a recent study identifying sex-specific piriform projections involved in emotion discrimination using a pain-based EDT paradigm^40^. Differences in emotional triggers (pain vs. mild restraint) and testing duration may explain the absence of sex-specific behavior in our task. Together, these studies suggest that sex-dependent circuit engagement in emotion recognition depends on both stimulus salience and temporal dynamics.

In humans, only male subjects showed amygdala activation in response to angry faces during the EFT, mirroring findings in our nEDT. Additionally, sex differences in prefrontal-amygdala connectivity emerged: males showed stronger vmPFC-amygdala coupling, while females exhibited increased OFC-amygdala connectivity. These results echo prior work linking cortico-amygdala connectivity to emotional reactivity and empathic responses in a sex-dependent manner^55–57^. Anatomical correspondence between rodent MO and human OFC subregions is imperfect. Indeed, rodents lack the granular OFC subdivisions found in humans^31^, making direct anatomical homology with the human vmPFC challenging. However, agranular regions in the mouse MO are often considered putative homologs of subregions from the human vmPFC/OFC, based on shared connectivity and functional roles in value-based decision-making and emotion regulation^58,59^. Thus, the results of our study strongly suggest the involvement of cortico-amygdala circuits in both species as a conserved mechanism for processing negative emotional states.

According to previous studies showing age and sex differences in emotion recognition^36^, our final set of findings revealed that aged male, but not female, mice lose the ability to discriminate emotional states. This behavioral decline was accompanied by reduced activity in BLA and MO, contrasting with their robust activation in young males. This aligns with studies linking MO damage to impaired emotion recognition^33^, although some human reports suggest that the amygdala may not directly mediate age-related deficits^37,60^. Importantly, chemogenetic activation of the BLA restored emotion recognition in aged males without affecting general behavior, providing causal evidence for its role in emotion processing and offering a potential therapeutic entry point.

Overall, our study identifies the BLA and its reciprocal connections with the MO as key regulators of emotion recognition in a sex- and age-dependent manner. By integrating behavioral, anatomical, photometry, chemogenetic, and human fMRI approaches, we demonstrate that cortico-amygdala circuits are dynamically and differentially recruited across sex and age. These findings suggest a complex network underlying emotion recognition and underscore the importance of sex and age as biological variables in socio-emotional neuroscience.

## Acknowledgements

We would like to thank the personnel of the Animal Facility of the Parc de Recerca Biomedica de Barcelona (PRBB) for mouse care. We thank all the members of our lab for useful discussions during the development of the project and Remi Proville (Aquineuro, Bordeaux, France) for the great help in the analysis of *in vivo* photometry. This work was supported by la Generalitat de Catalunya (SGR (00022); the European Research Council (ERC) under the European Union’s Horizon 2020 research and innovation programme (Grant agreement No. [948217]); the BBVA Foundation (“Proyecto realizado con la Beca Leonardo a Investigadores y Creadores Culturales 2024 de la Fundación BBVA”, LEO24-1-12486) and the Ministry of Economy and Competitiveness (ERASE project).

## Author Contributions

A.B-G. and JA.G-P. conceived the project. A.B-G., X.C. and JA.G-P designed the experiments. JA.G-P, I.P.A-J, P.S-C., V.A. performed the experiments. JA.G-P, I.P.A-J. M.C-G., L.T. and B.D. analyzed the data with input from A.B-G., X.C., F.P and G.S. I.P.A-J. contributed to the development of analytical tools. L.V.P helped with specific behavioral experiments. A.B-G., I.A.J, X.C. and JA.G-P interpreted the results. A.B-G. and JA.G-P. wrote the manuscript with input from all authors. All authors discussed the results and approved the final version of the manuscript.

## References

1 Beaudoin, C. & Beauchamp, M. H. Social cognition. Handb Clin Neurol 173, 255–264 (2020). 10.1016/B978-0-444-64150-2.00022-8

2 Wascher, C. A. F., Kulahci, I. G., Langley, E. J. G. & Shaw, R. C. How does cognition shape social relationships? Philos Trans R Soc Lond B Biol Sci 373 (2018). 10.1098/rstb.2017.0293

3 Connolly, H. L., Lefevre, C. E., Young, A. W. & Lewis, G. J. Emotion recognition ability: Evidence for a supramodal factor and its links to social cognition. Cognition 197, 104166 (2020). 10.1016/j.cognition.2019.104166

4 Masoomi, M. et al. Emotion recognition deficits in children and adolescents with autism spectrum disorder: a comprehensive meta-analysis of accuracy and response time. Front Child Adolesc Psychiatry 3, 1520854 (2024). 10.3389/frcha.2024.1520854

5 Kohler, C. G., Turner, T. H., Gur, R. E. & Gur, R. C. Recognition of facial emotions in neuropsychiatric disorders. CNS Spectr 9, 267–274 (2004). 10.1017/s1092852900009202

6 Sola, C. et al. Understanding basic and social emotions in Alzheimer’s disease and frontotemporal dementia. Front Psychol 16, 1535722 (2025). 10.3389/fpsyg.2025.1535722

7 Rafiee, Y. & Schacht, A. Sex differences in emotion recognition: investigating the moderating effects of stimulus features. Cogn Emot 37, 863–873 (2023). 10.1080/02699931.2023.2222579

8 Maltese, F. et al. Self-experience of a negative event alters responses to others in similar states through prefrontal cortex CRF mechanisms. Nat Neurosci 28, 122–136 (2025). 10.1038/s41593-024-01816-y

9 Cortes, D. S., Tornberg, C., Banziger, T., Elfenbein, H. A., Fischer, H. & Laukka, P. Effects of aging on emotion recognition from dynamic multimodal expressions and vocalizations. Sci Rep 11, 2647 (2021). 10.1038/s41598-021-82135-1

10 Langford, D. J. et al. Social approach to pain in laboratory mice. Soc Neurosci 5, 163–170 (2010). 10.1080/17470910903216609

11 Dolensek, N., Gehrlach, D. A., Klein, A. S. & Gogolla, N. Facial expressions of emotion states and their neuronal correlates in mice. Science 368, 89–94 (2020). 10.1126/science.aaz9468

12 Scheggia, D. et al. Somatostatin interneurons in the prefrontal cortex control affective state discrimination in mice. Nat Neurosci 23, 47–60 (2020). 10.1038/s41593-019-0551-8

13 Ferretti, V. et al. Oxytocin Signaling in the Central Amygdala Modulates Emotion Discrimination in Mice. Curr Biol 29, 1938–1953 e1936 (2019). 10.1016/j.cub.2019.04.070

14 Dautan, D. et al. Cortico-cortical transfer of socially derived information gates emotion recognition. Nat Neurosci 27, 1318–1332 (2024). 10.1038/s41593-024-01647-x

15 Keysers, C., Knapska, E., Moita, M. A. & Gazzola, V. Emotional contagion and prosocial behavior in rodents. Trends Cogn Sci 26, 688–706 (2022). 10.1016/j.tics.2022.05.005

16 Gachomba, M. J. M., Esteve-Agraz, J. & Marquez, C. Prosocial behaviors in rodents. Neurosci Biobehav Rev 163, 105776 (2024). 10.1016/j.neubiorev.2024.105776

17 Dal Bo, E., et al. Emotion perception through the nose: how olfactory emotional cues modulate the perception of neutral facial expressions in affective disorders. Transl Psychiatry 14, 342 (2024). 10.1038/s41398-024-03038-z

18 Allsop, S. A. et al. Corticoamygdala Transfer of Socially Derived Information Gates Observational Learning. Cell 173, 1329–1342 e1318 (2018). 10.1016/j.cell.2018.04.004

19 Terranova, J. I. et al. Hippocampal-amygdala memory circuits govern experience-dependent observational fear. Neuron 110, 1416–1431 e1413 (2022). 10.1016/j.neuron.2022.01.019

20 Scheggia, D. et al. Reciprocal cortico-amygdala connections regulate prosocial and selfish choices in mice. Nat Neurosci 25, 1505–1518 (2022). 10.1038/s41593-022-01179-2

21 Sun, W. et al. Reviving-like prosocial behavior in response to unconscious or dead conspecifics in rodents. Science 387, eadq2677 (2025). 10.1126/science.adq2677

22 Wu, Y. et al. Sex-specific neural circuits of emotion regulation in the centromedial amygdala. Sci Rep 6, 23112 (2016). 10.1038/srep23112

23 DeNardo, L. A. et al. Temporal evolution of cortical ensembles promoting remote memory retrieval. Nat Neurosci 22, 460–469 (2019). 10.1038/s41593-018-0318-7

24 Gonzalez-Parra, J. A., Acciai, V., Vidal-Palencia, L., Canela-Grimau, M. & Busquets-Garcia, A. Projecting neurons from the lateral entorhinal cortex to the basolateral amygdala mediate the encoding of incidental odor-taste associations. Proc Natl Acad Sci U S A 122, e2502127122 (2025). 10.1073/pnas.2502127122

25 Mathis, A. et al. DeepLabCut: markerless pose estimation of user-defined body parts with deep learning. Nat Neurosci 21, 1281–1289 (2018). 10.1038/s41593-018-0209-y

26 Weinreb, C. et al. Keypoint-MoSeq: parsing behavior by linking point tracking to pose dynamics. Nat Methods 21, 1329–1339 (2024). 10.1038/s41592-024-02318-2

27 Canela-Grimau, M., Pinho, J. S. & Busquets-Garcia, A. Profiling mouse behavior with computational tools to assess age-dependent differences in associative learning. Cell Rep Methods 5, 101144 (2025). 10.1016/j.crmeth.2025.101144

28 Schumann, G. et al. The IMAGEN study: reinforcement-related behaviour in normal brain function and psychopathology. Mol Psychiatry 15, 1128–1139 (2010). 10.1038/mp.2010.4

29 Grosbras, M. H. & Paus, T. Brain networks involved in viewing angry hands or faces. Cereb Cortex 16, 1087–1096 (2006). 10.1093/cercor/bhj050

30 Rolls, E. T., Huang, C. C., Lin, C. P., Feng, J. & Joliot, M. Automated anatomical labelling atlas 3. Neuroimage 206, 116189 (2020). 10.1016/j.neuroimage.2019.116189

31 Preuss, T. M. & Wise, S. P. Evolution of prefrontal cortex. Neuropsychopharmacology 47, 3–19 (2022). 10.1038/s41386-021-01076-5

32 Mumford, J. A. & Ramsey, J. D. Bayesian networks for fMRI: a primer. Neuroimage 86, 573–582 (2014). 10.1016/j.neuroimage.2013.10.020

33 Nakajima, R., Kinoshita, M., Okita, H. & Nakada, M. Posterior-prefrontal and medial orbitofrontal regions play crucial roles in happiness and sadness recognition. Neuroimage Clin 35, 103072 (2022). 10.1016/j.nicl.2022.103072

34 Jabarin, R., Mohapatra, A. N., Ray, N., Netser, S. & Wagner, S. Distinct prelimbic cortex neuronal responses to emotional states of others drive emotion recognition in adult mice. Curr Biol 35, 994–1011 e1018 (2025). 10.1016/j.cub.2025.01.014

35 Fujima, S., Sato, M., Nakai, N. & Takumi, T. Parvalbumin interneurons in the insular cortex control social familiarity and emotion recognition. Cell Rep 44, 116085 (2025). 10.1016/j.celrep.2025.116085

36 Abbruzzese, L., Magnani, N., Robertson, I. H. & Mancuso, M. Age and Gender Differences in Emotion Recognition. Front Psychol 10, 2371 (2019). 10.3389/fpsyg.2019.02371

37 Malykhin, N., Pietrasik, W., Aghamohammadi-Sereshki, A., Ngan Hoang, K., Fujiwara, E. & Olsen, F. Emotional recognition across the adult lifespan: Effects of age, sex, cognitive empathy, alexithymia traits, and amygdala subnuclei volumes. J Neurosci Res 101, 367–383 (2023). 10.1002/jnr.25152

38 Rogers-Carter, M. M., Djerdjaj, A., Culp, A. R., Elbaz, J. A. & Christianson, J. P. Familiarity modulates social approach toward stressed conspecifics in female rats. PLoS One 13, e0200971 (2018). 10.1371/journal.pone.0200971

39 Langford, D. J. et al. Varying perceived social threat modulates pain behavior in male mice. J Pain 12, 125–132 (2011). 10.1016/j.jpain.2010.06.003

40 Fang, S. et al. Sexually dimorphic control of affective state processing and empathic behaviors. Neuron 112, 1498–1517 e1498 (2024). 10.1016/j.neuron.2024.02.001

41 Misiolek, K. et al. Prosocial behavior, social reward and affective state discrimination in adult male and female mice. Sci Rep 13, 5583 (2023). 10.1038/s41598-023-32682-6

42 Dilli Batcha, J. S., et al. Factors Influencing Pharmacokinetics of Tamoxifen in Breast Cancer Patients: A Systematic Review of Population Pharmacokinetic Models. Biology (Basel) 12 (2022). 10.3390/biology12010051

43 Parel, S. T. & Pena, C. J. Genome-wide Signatures of Early-Life Stress: Influence of Sex. Biol Psychiatry 91, 36–42 (2022). 10.1016/j.biopsych.2020.12.010

44 Hernandez-Vivanco, A., de la Vega-Ruiz, R., Montes-Mellado, A., Azcoitia, I. & Mendez, P. Activational and organizational effects of sex hormones on hippocampal inhibitory neurons. J Neurosci 45 (2025). 10.1523/JNEUROSCI.1764-24.2025

45 Ossewaarde, L. et al. Neural mechanisms underlying changes in stress-sensitivity across the menstrual cycle. Psychoneuroendocrinology 35, 47–55 (2010). 10.1016/j.psyneuen.2009.08.011

46 Shafir, T. et al. Postmenopausal hormone use impact on emotion processing circuitry. Behav Brain Res 226, 147–153 (2012). 10.1016/j.bbr.2011.09.012

47 Park, S., Cho, J. & Huh, Y. Role of the anterior insular cortex in restraint-stress induced fear behaviors. Sci Rep 12, 6504 (2022). 10.1038/s41598-022-10345-2

48 Schneider, S. et al. Boys do it the right way: sex-dependent amygdala lateralization during face processing in adolescents. Neuroimage 56, 1847–1853 (2011). 10.1016/j.neuroimage.2011.02.019

49 Frith, C. D. & Frith, U. Mechanisms of social cognition. Annu Rev Psychol 63, 287–313 (2012). 10.1146/annurev-psych-120710-100449

50 Shi, W., Meisner, O. C., Blackmore, S., Jadi, M. P., Nandy, A. S. & Chang, S. W. C. The orbitofrontal cortex: A goal-directed cognitive map framework for social and non-social behaviors. Neurobiol Learn Mem 203, 107793 (2023). 10.1016/j.nlm.2023.107793

51 Calder, A. J., Keane, J., Manes, F., Antoun, N. & Young, A. W. Impaired recognition and experience of disgust following brain injury. Nat Neurosci 3, 1077–1078 (2000). 10.1038/80586

52 Vieitas-Gaspar, N., Soares-Cunha, C. & Rodrigues, A. J. From valence encoding to motivated behavior: A focus on the nucleus accumbens circuitry. Neurosci Biobehav Rev 172, 106125 (2025). 10.1016/j.neubiorev.2025.106125

53 Song, Z., Swarna, S. & Manns, J. R. Prioritization of social information by the basolateral amygdala in rats. Neurobiol Learn Mem 184, 107489 (2021). 10.1016/j.nlm.2021.107489

54 Chang, C. H. & Grace, A. A. Inhibitory Modulation of Orbitofrontal Cortex on Medial Prefrontal Cortex-Amygdala Information Flow. Cereb Cortex 28, 1–8 (2018). 10.1093/cercor/bhw342

55 Liang, X., Zebrowitz, L. A. & Aharon, I. Effective connectivity between amygdala and orbitofrontal cortex differentiates the perception of facial expressions. Soc Neurosci 4, 185–196 (2009). 10.1080/17470910802453105

56 Domes, G. et al. The neural correlates of sex differences in emotional reactivity and emotion regulation. Hum Brain Mapp 31, 758–769 (2010). 10.1002/hbm.20903

57 Rodriguez-Nieto, G., Mercadillo, R. E., Pasaye, E. H. & Barrios, F. A. Affective and cognitive brain-networks are differently integrated in women and men while experiencing compassion. Front Psychol 13, 992935 (2022). 10.3389/fpsyg.2022.992935

58 Myers-Schulz, B. & Koenigs, M. Functional anatomy of ventromedial prefrontal cortex: implications for mood and anxiety disorders. Mol Psychiatry 17, 132–141 (2012). 10.1038/mp.2011.88

59 Rolls, E. T., Cheng, W. & Feng, J. The orbitofrontal cortex: reward, emotion and depression. Brain Commun 2, fcaa196 (2020). 10.1093/braincomms/fcaa196

60 Orlando, I., Ricci, C., Griffanti, L. & Filippini, N. Neural correlates of successful emotion recognition in healthy elderly: a multimodal imaging study. Soc Cogn Affect Neurosci 18 (2023). 10.1093/scan/nsad058

