## Supplementary Material for "Sex-dependent cortico-amygdala circuits controlling emotion recognition"

**Correspoding author:**

Arnau Busquets-Garcia, PhD
Cell-Type Mechanisms in Normal and Pathological Behavior Research Group
Neuroscience Research Program, Hospital del Mar Research Institute.
PRBB
Calle Dr. Aiguader 88
Barcelona 08003
SPAIN
Mail:

**Content:**

**Supplementary Figures 1-4**

**Supplementary Tables 1-2**

#### Supplementary Figures

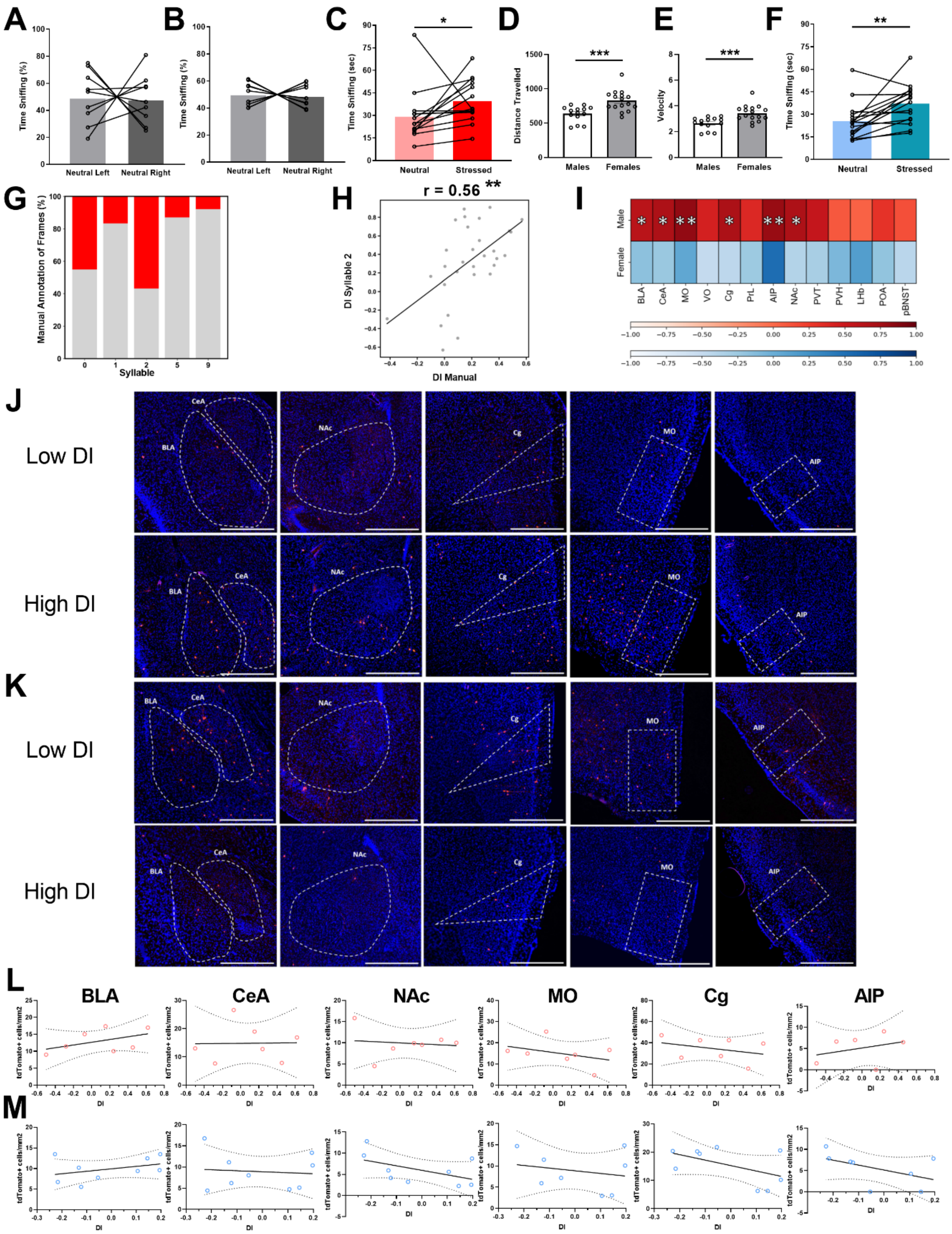


**Supplementary Figure 1. Behavioral performance of young TRAP2:Ai14 mice in a negative Affective Discrimination Task.** (A-B) Percentage of time spent sniffing the neutral compartment on the left versus right side during the task in control young male (A) and female (B) TRAP2:Ai14 mice. (C) Raw time (seconds) spent sniffing neutral versus stressed demonstrators in male TRAP2:Ai14 mice. (D–E) Comparison of distance traveled (D) and velocity (E) between male and female TRAP2:Ai14 mice in the EDT group. (F) Raw time (seconds) spent sniffing neutral versus stressed demonstrators in female TRAP2:Ai14 mice. (G) Percentage of manually annotated sniffing time (red) versus other behaviors (gray) across individual behavioral syllables. (H) Pearson correlation between the DI calculated from manually annotated sniffing time and DI in syllable 2. (I) Correlation analyses between tdTomato+ cells per $mm^2$ and the DI of sniffing time in young male and female TRAP2:Ai14 mice across all brain regions analyzed. (J-K) Representative images of tdTomato expression in the BLA, CeA, NAc, Cg, MO, and AIP in High DI and Low DI male (J) and female (K) TRAP2:Ai14 mice from the EDT group.(L) Correlation analyses between tdTomato+ cells per $mm^2$ and DI (time spent sniffing neutral right vs. neutral left) in Control group young male TRAP2:Ai14 mice for the BLA, CeA, NAc, MO, Cg, and AIP. (M) Equivalent correlation analyses in Control group young female TRAP2:Ai14 mice for the BLA, CeA, NAc, MO, Cg, and AIP. BLA: Basolateral Amygdala; CeA: Central Amygdala; NAc: Nucleus Accumbens; Cg: Cingulate cortex; MO: Medial Orbitofrontal cortex; AIP: Agranular Posterior Insular cortex; PVT: Paraventricular nucleus of the Thalamus; PVH: Paraventricular nucleus of the Hypothalamus; POA: Preoptic area; pBNST: posterior Bed Nucleus of the Stria Terminalis. Fluorescent images represent 10x of tdTomato cells. (Scale bar, 500 μm). Data are represented as before-after for the percentage of time sniffing or as mean ± SEM for locomotion. In correlation analysis, lines represent linear regression fits; shaded areas show 95% confidence intervals. ***$P < 0.001$; **$P < 0.01$; *$P < 0.05$. For statistical details, see SI Appendix, Table S2.

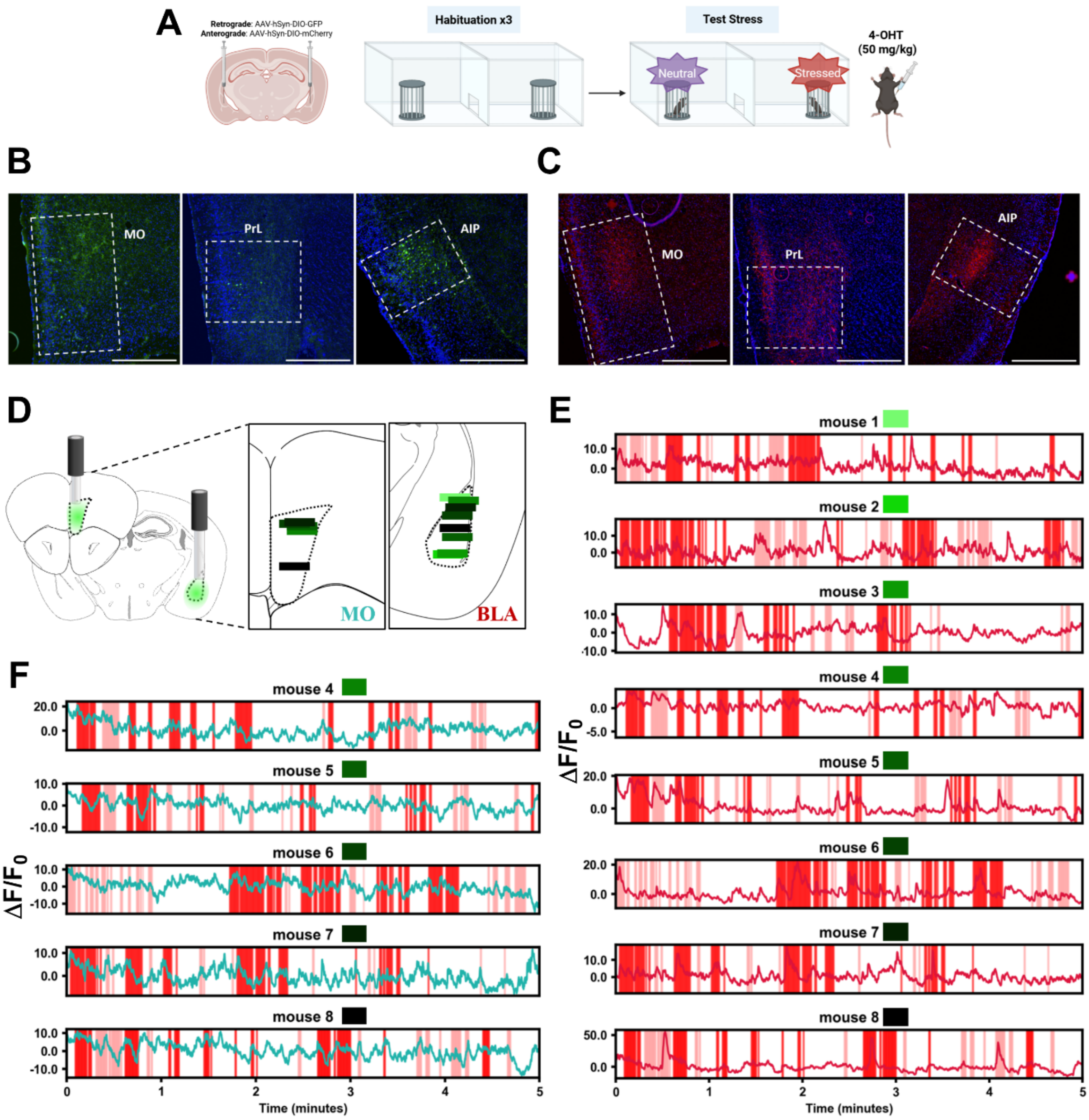


**Supplementary Figure 2. Dynamic BLA and MO engagement during negative emotion recognition.** (A) Schematic representation of bilateral infusions of Cre-dependent anterograde tracers (AAV-hSyn-DIO-mCherry) or Cre-dependent retrograde tracers (rAAV-hSyn-DIO-GFP) into the BLA of TRAP2:Ai14 mice, used to map activity-dependent outputs from and inputs into the BLA during emotion recognition. (B–C) Representative images of BLA inputs (retrograde, GFP) from the MO, PrL, and AIP (B) and BLA outputs (anterograde, mCherry) to the MO, PrL, and AIP (C). (D) Schematic showing surgery set up (left) and placements of tip of the optic fiber in MO (middle) and BLA (right). (E) BLA recorded fluorescence as $\Delta F/F_0$ for all analyzed mice for the first 4 minutes of the test, where downstream analysis was carried. Colored patches indicate the time periods in which mice sniffed the stressed demonstrator (red) or the neutral (pink) (F) Equivalent as E, for MO. Fluorescent images represent 10x of mCherry terminals or GFP cells. (Scale bar, 500 μm).

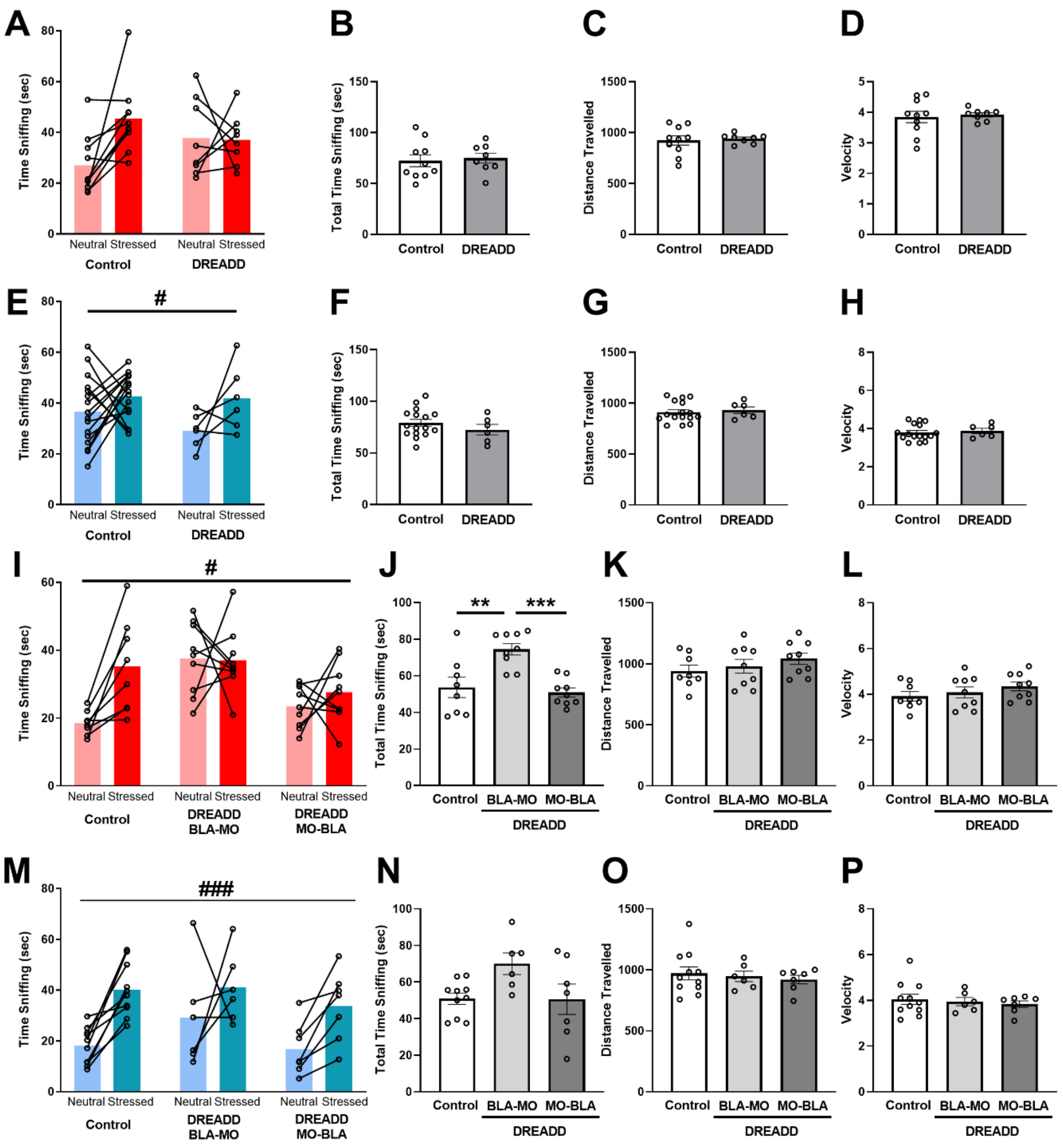


**Supplementary Figure 3. Chemogenetic inhibition of the BLA and reciprocal BLA–MO projections during emotion recognition.** (A) Raw time (seconds) spent sniffing neutral versus stressed demonstrators in control and DREADD male groups. (B) Total sniffing time (seconds) directed toward neutral and stressed demonstrators in control and DREADD male groups. (C-D) Comparison of distance traveled (C) and velocity (D) between control and DREADD male mice. (E) Raw time (seconds) spent sniffing neutral versus stressed demonstrators in control and DREADD female groups. (F) Total sniffing time (seconds) directed toward neutral and stressed demonstrators in control and DREADD female groups. (GH) Comparison of distance traveled (G) and velocity (H) between control and DREADD female mice. (I) Raw time (seconds) spent sniffing neutral versus stressed demonstrators in control, BLA MO, and MO-BLA male groups. (J) Total sniffing time (seconds) directed toward neutral and stressed demonstrators in control, BLA-MO, and MO-BLA male groups. (K-L) Comparison of distance traveled (K) and velocity (L) between control, BLA-MO, and MO BLA male mice. (M) Raw time (seconds) spent sniffing neutral versus stressed demonstrators in control, BLA-MO, and MO-BLA female groups. (N) Total sniffing time (seconds) directed toward neutral and stressed demonstrators in control, BLA-MO, and MO-BLA female groups. (O-P) Comparison of distance traveled (O) and velocity (P) between control, BLA-MO, and MO-BLA female mice. Data are represented as before-after for the percentage of time sniffing or as mean ± SEM for locomotion. # Denote significant effect of demonstrator factor (neutral versus stressed). ***/### $P < 0.001$; **/## $P < 0.01$; */# $P < 0.05$. For statistical details, see SI Appendix, Table S2.

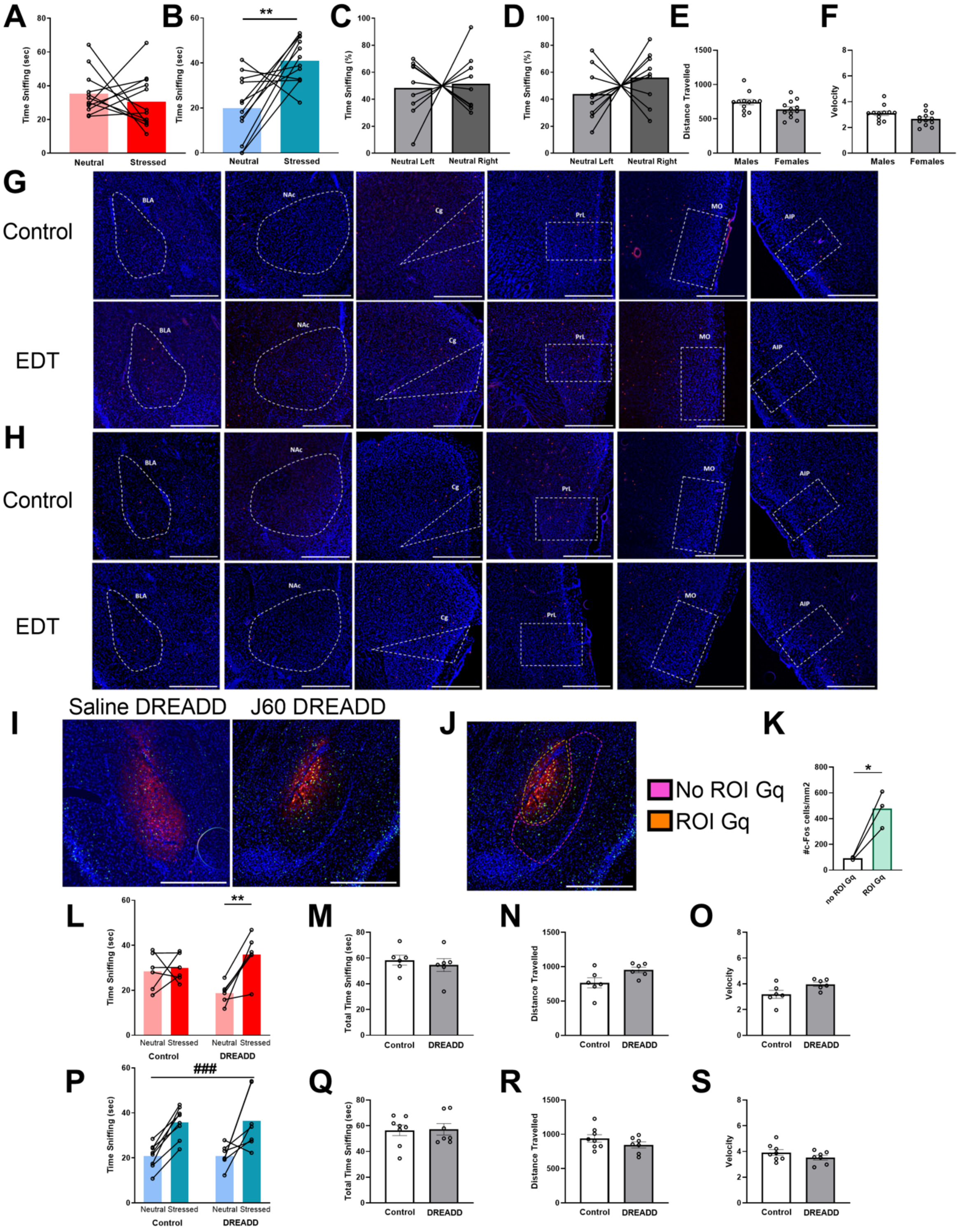


**Supplementary Figure 4. Age-dependent involvement of amygdala in emotion recognition.** (A-B) Raw time (seconds) spent sniffing neutral versus stressed demonstrators in aged male (A) and female (B) TRAP2:Ai14 mice. (C-D) Percentage of time spent sniffing the neutral compartment on the left versus right side in control aged male (C) and female (D) TRAP2:Ai14 mice. (E–F) Comparison of distance traveled (E) and velocity (F) between aged male and female TRAP2:Ai14 mice in the EDT group. (G–H) Representative images of tdTomato expression in the BLA, NAc, Cg, MO, and AIP in control and EDT aged male (G) and female (H) TRAP2:Ai14 groups. (I) Representative BLA images of excitatory DREADD-mCherry expression (red) and c-Fos+ cells (green) in animals injected with saline (left) or J60 (right). (J) Representative BLA image depicting the ROIs used for c-Fos analysis, showing an ROI with excitatory DREADD expression (orange) and a no-ROI control area lacking expression (pink). (K) Quantification of c-Fos+ cells per $mm^2$ in ROI-Gq versus no ROI-Gq areas in animals injected with J60. (L) Raw time (seconds) spent sniffing neutral versus stressed demonstrators in control and DREADD aged male groups. (M) Total sniffing time (seconds) directed toward neutral and stressed demonstrators in control and DREADD aged male groups. (N-O) Comparison of distance traveled (N) and velocity (O) between control and DREADD aged male mice. (P) Raw time (seconds) spent sniffing neutral versus stressed demonstrators in control and DREADD aged female groups. (Q) Total sniffing time (seconds) directed toward neutral and stressed demonstrators in control and DREADD aged female groups. (R-S) Comparison of distance traveled (R) and velocity (S) between control and DREADD aged female mice. Fluorescent images represent 10x of tdTomato+ cells, mCherry signal expression and c-Fos+ cells. (Scale bar, 500 μm). Data are represented as before-after for the percentage of time sniffing or as mean ± SEM for the locomotion and quantification of c-Fos+ cells. In the repeated measure two-way ANOVA statistical analysis * denote significant interaction between the two factors while # denote significant effect of demonstrator factor (neutral versus stressed). ***/### $P < 0.001$; **/## $P < 0.01$; */# $P < 0.05$. For statistical details, see SI Appendix, Table S1.

### Table S1. Statistical analysis. Related to Main Figures 1-5

| *Experiment* | *Sample size per group* | *Passed normality test?* | *Statistical test* | *Statistical value* |
|---|---|---|---|---|
| C. TRAP2 young males EDT group. Time sniffing (%) | n=14 | Yes | Paired t test/One sample t test | **t=2,664, df=13, p<0.05** |
| D. TRAP2 young males EDT group. Time in ROI (%) | n=14 | Yes | Paired t test/One sample t test | **t=3,116, df=13, p<0.01** |
| F. TRAP2 young females EDT group. Time sniffing (%) | n=16 | Yes | Paired t test/One sample t test | **t=4,313, df=15, p<0.001** |
| G. TRAP2 young females EDT group. Time in ROI (%) | n=16 | Yes | Paired t test/One sample t test | **t=1,781, df=15, p= n.s** |
| I. TRAP2 young males EDT group. Syllable 2 (%) | n=14 | Yes | Paired t test/One sample t test | **t=3,089, df=13, p<0.01** |
| J. TRAP2 young females EDT group. Syllable 2 (%) | n=16 | Yes | Paired t test/One sample t test | **t=2,976, df=15, p<0.01** |
| K. Correlation EDT group BLA males | n=11 | Yes | Pearson Correlation | **r=0,6101, p<0.05** |
| L. Correlation EDT group CeA males | n=11 | Yes | Pearson Correlation | **r=0,6949; p<0.05** |
| M. Correlation EDT group NAc males | n=11 | Yes | Pearson Correlation | **r=0,6895; p<0.05** |
| N. Correlation EDT group MO males | n=11 | Yes | Pearson Correlation | **r=0,8233; p<0.01** |
| O. Correlation EDT group Cg males | n=12 | Yes | Pearson Correlation | **r=0,703, p<0.05** |
| P. Correlation EDT group AIP males | n=9 | Yes | Pearson Correlation | **r=0,8007, p<0.01** |
| Q. Correlation EDT group BLA females | n=14 | Yes | Pearson Correlation | **r=-0,1712, p=n.s** |
| R. Correlation EDT group CeA females | n=14 | Yes | Pearson Correlation | **r=-0,2329, p=n.s** |
| S. Correlation EDT group NAc females | n=13 | Yes | Pearson Correlation | **r=-0,3771, p=n.s** |
| T. Correlation EDT group MO females | n=13 | Yes | Pearson Correlation | **r=0,06687, p=n.s** |
| U. Correlation EDT group Cg females | n=14 | Yes | Pearson Correlation | **r=-0,439, p=n.s** |
| W. Correlation EDT group AIP females | n=10 | Yes | Pearson Correlation | **r=0,5305, p=n.s** |
| B. Time sniffing (%) | n=8 | Yes | Paired t test | **t=5,410, df=7, p<0.001** |
| F. Sniffing vs other behavior BLA | n = 8 | Yes | Paired t test/One sample t test | **t=5.268, df=7, p<0.01** |
| F. Sniffing vs other behavior MO | n = 5 | Yes | Paired t test/One sample t test | **t=3.918, df=4, p<0.05** |
| I,L. Sniffing onset/offset BLA | n = 8 | NA | Bootstrap 95% CI | **Outside 95% CI** |
| I,L.Sniffing onset/offset MO | n = 5 | NA | Bootstrap 95% CI | **Outside 95% CI** |
| C. BLA inhibition males (%) | n=8-10 | Yes | 2 way ANOVA repeated measures (Bonferroni) | **Interaction: F (1, 16) = 4,883, p<0.05** |
| | | | | **Neutral vs Stressed (Control) p<0.01** |
| D. BLA inhibition females (%) | n=6-16 | Yes | 2 way ANOVA repeated measures | **Manipulation effect: F (1, 20) = 1795, p<0.001** |
| | | | | **Demonstrator effect: F (1, 20) = 4,895, p<0.05** |
| I. BLA-MO/MO-BLA inhibition males (%) | n=8-9 | Yes | 2 way ANOVA repeated measures (Bonferroni) | **Interaction: F (2, 23) = 3,505, p<0.05** |
| | | | | **Neutral vs Stressed (Control) p<0.01** |
| J. BLA-MO/MO-BLA inhibition males (%) | n=6-10 | Yes | 2 way ANOVA repeated measures | **Demonstrator effect: F (1, 20) = 28,60, p<0.001** |
| B. Male Amygdala Neutral vs Angry | n=1340 | Yes | Paired t test | **t = 5.24, p<0.05 FDR correction** |
| B. Male vmPFC Neutral vs Angry | n=1340 | Yes | Paired t test | **t = 9.30, p<0.05 FDR correction** |
| B. Male postOFC Neutral vs Angry | n=1340 | Yes | Paired t test | **t = 3.88, p<0.05 FDR correction** |
| B. Female vmPFC Neutral vs Angry | n=1340 | Yes | Paired t test | **t = 11.56, p<0.05 FDR correction** |
| C. Male Amygdala-vmPFC connectivity | n=1340 | Yes | One sample t test | **p<0.001** |
| C. Female Amygdala-OFC connectivity | n=1340 | Yes | One sample t test | **p<0.001** |
| B. TRAP2 aged males ADT group. Time sniffing (%) | n=12 | Yes | Paired t test/One sample t test | **t=0,9203, df=11, p=n.s** |
| C. TRAP2 aged females ADT group. Time sniffing (%) | n=12 | Yes | Paired t test/One sample t test | **t=3,276, df=11, p<0.01** |
| D. BLA aged males | n=7-11 | Yes | Unpaired t test | **t=0,2287, df=16, p=n.s** |
| E. MO aged males | n=7-10 | Yes | Unpaired t test | **t=1,354, df=16, p=n.s** |
| F. BLA aged females | n=8-10 | No | Mann-Whitney Test | **p=n.s** |
| G. MO aged females | n=8-10 | Yes | Unpaired t test | **t=1,354, df=16, p=n.s** |
| H. BLA aged males Correlation | n=11 | Yes | Pearson Correlation | **r=0,0984, p=n.s** |
| I. MO aged males Correlation | n=11 | Yes | Pearson Correlation | **r=-0,0656, p=n.s** |
| J. BLA aged females Correlation | n=10 | Yes | Pearson Correlation | **r=-0,121, p=n.s** |
| K. MO aged females Correlation | n=10 | Yes | Pearson Correlation | **r=-0,2707, p=n.s** |
| M. BLA activation males (%) | n=6 | Yes | 2 way ANOVA repeated measures (Bonferroni) | **Interaction: F (1, 10) = 7,159** |
| | | | | **Neutral vs Stressed (DREADD) p<0.01** |
| N. BLA activation females (%) | n=7-8 | Yes | 2 way ANOVA repeated measures | **Mnipulation effect: F (1, 13) = 17,76, p<0.01** |
| | | | | **Demonstrator effect: F (1, 13) = 35,00, p<0.001** |

### Table S2. Statistical analysis. Related to Supplemental Figures 1-4

| *Figure* | *Experiment* | *Sample size per group* | *Passed normality test?* | *Statistical test* | *Statistical value* |
|---|---|---|---|---|---|
| 1 | A. TRAP2 young males Control group. Time sniffing (%) | n=9 | No | Wilcoxon Test | **t=0,1279, df=8, p=n.s** |
| | B. TRAP2 young female Control group. Time sniffing (%) | n=9 | No | Wilcoxon Test | **t=0,2112, df=8, p=n.s** |
| | C. TRAP2 young males EDT group. Time sniffing (sec) | n=14 | No | Wilcoxon Test | **p<0.05** |
| | D. Distance Travelled male vs female | n=14-16 | No | Mann-Whitney Test | **p<0.001** |
| | E. Velocity male vs female | n=14-16 | No | Mann-Whitney Test | **p<0.001** |
| | F. TRAP2 young females EDT group. Time sniffing (sec) | n=16 | No | Wilcoxon Test | **p<0.01** |
| | H. Correlation DI manual vs DI syllable 2 | n=30 | Yes | Pearson Correlation | **r=0.56, p<0.01** |
| | L. Correlation Control group BLA males | n=7 | Yes | Pearson Correlation | **r=0,4723, p=n.s** |
| | L. Correlation Control group CeA males | n=7 | Yes | Pearson Correlation | **r=0,01845, p=n.s** |
| | L. Correlation Control group NAc males | n=7 | Yes | Pearson Correlation | **r=-0,1239, p=n.s** |
| | L. Correlation Control group MO males | n=7 | Yes | Pearson Correlation | **r=-0,3789, p=n.s** |
| | L. Correlation Control group Cg males | n=7 | Yes | Pearson Correlation | **r=-0,3335, p=n.s** |
| | L. Correlation Control group AIP males | n=6 | Yes | Pearson Correlation | **r=0,3234, p=n.s** |
| | M. Correlation Control group BLA females | n=9 | Yes | Pearson Correlation | **r=0,3602, p=n.s** |
| | M. Correlation Control group CeA females | n=9 | Yes | Pearson Correlation | **r=-0,09345, p=n.s** |
| | M. Correlation Control group NAc females | n=9 | Yes | Pearson Correlation | **r=-0,5180, p=n.s** |
| | M. Correlation Control group MO females | n=8 | Yes | Pearson Correlation | **r=-0,2201, p=n.s** |
| | M. Correlation Control group Cg females | n=9 | No | Spearman Correlation | **r=-0,2833, p=n.s** |
| | M. Correlation Control group AIP females | n=8 | Yes | Pearson Correlation | **r=-0,4983, p=n.s** |
| 3 | A. BLA inhibition males (sec) | n=8-10 | Yes | 2 way ANOVA repeated measures | **Interaction: F (1, 16) = 4,150, p=0.058 n.s** |
| | B. Total time sniffing male Control vs DREADD | n=8-10 | Yes | Unpaired t Test | **t=0,3154, df=16, p=n.s** |
| | C. Distance Travelled male Control vs DREADD | n=8-10 | Yes | Unpaired t Test | **t=0,3326, df=16, p=n.s** |
| | D. Velocity male Control vs DREADD | n=8-10 | Yes | Unpaired t Test | **t=0,3326, df=16, p=n.s** |
| | E. BLA inhibition females (sec) | n=6-16 | Yes | 2 way ANOVA repeated measures | **Demonstrator effect: F (1, 20) = 4,465, p<0.05** |
| | F. Total time sniffing female Control vs DREADD | n=6-16 | Yes | Unpaired t Test | **t=1,042, df=20, p=n.s** |
| | G. Distance Travelled female Control vs DREADD | n=6-16 | Yes | Unpaired t Test | **t=0,4271, df=20, p=n.s** |
| | H. Velocity female Control vs DREADD | n=6-16 | Yes | Unpaired t Test | **t=0,4271, df=20, p=n.s** |
| | I. BLA-MO/MO-BLA inhibition male (sec) | n=8-9 | Yes | 2 way ANOVA repeated measures | **Manipulation effect:F (2, 23) = 11,73, p<0.001**<br>**Demonstrator effect: F (1, 23) = 5,208, p<0.05** |
| | J. Total time sniffing male BLA-MO/MO-BLA | n=8-9 | Yes | 1 way ANOVA | **F (2, 23) = 11,73, p<0.001** |
| | K. Distance Travelled male BLA-MO/MO-BLA | n=8-9 | Yes | 1 way ANOVA | **F (2, 23) = 0,9964, p=n.s** |
| | L. Velocity male BLA-MO/MO-BLA | n=8-9 | Yes | 1 way ANOVA | **F (2, 23) = 0,9964, p=n.s** |
| | M. BLA-MO/MO-BLA inhibition female (sec) | n=6-10 | Yes | 2 way ANOVA repeated measures | **F (1, 20) = 18,37, p<0.001** |
| | N. Total time sniffing female BLA-MO/MO-BLA | n=6-10 | Yes | 1 way ANOVA | **F (2, 20) = 3,417, p= n.s** |
| | O. Distance Travelled female BLA-MO/MO-BLA | n=6-10 | Yes | 1 way ANOVA | **F (2, 21) = 0,2783, p=n.s** |
| | P. Velocity female BLA-MO/MO-BLA | n=6-10 | Yes | 1 way ANOVA | **F (2, 21) = 0,2783, p=n.s** |
| 4 | A. TRAP2 aged males EDT group. Time sniffing (sec) | n=12 | Yes | Paired t test | **t=0,6715, df=11, p=n.s** |
| | B. TRAP2 aged females EDT group. Time sniffing (sec) | n=12 | Yes | Paired t test | **t=3,484, df=11, p<0.01** |
| | C. TRAP2 young males Control group. Time sniffing (%) | n=9 | Yes | Paired t test | **t=0,2221, df=8, p=n.s** |
| | D. TRAP2 young female Control group. Time sniffing (%) | n=9 | Yes | Paired t test | **t=0,9348, df=8, p=n.s** |
| | E. Distance Travelled males vs females | n=12 | Yes | Unpaired t test | **t=1,911, df=22, p=n.s** |
| | F. Velocity males vs females | n=12 | Yes | Unpaired t test | **t=1,911, df=22, p=n.s** |
| | K. c-Fos Gq DREADD ROI vs noROI | n=3 | Yes | Paired t test | **t=5,048, df=2, p<0.05** |
| | L. BLA activation males (sec) | n=6 | Yes | 2 way ANOVA repeated measures (Bonferroni) | **Interaction: F (1, 10) = 7,597**<br>**Neutral vs Stressed (DREADD) p<0.01** |
| | M. Total time sniffing male Control vs DREADD | n=6 | Yes | Unpaired t Test | **t=0,6076, df=10, p=n.s** |
| | N. Distance Travelled male Control vs DREADD | n=6 | Yes | Unpaired t Test | **t=2,223, df=10, p=n.s** |
| | O. Velocity male Control vs DREADD | n=6 | Yes | Unpaired t Test | **t=2,223, df=10, p=n.s** |
| | P. BLA activation females (sec) | n=7-8 | Yes | 2 way ANOVA repeated measures | **Demonstrator effect: F (1, 13) = 28,12, p<0.001** |
| | Q. Total time sniffing female Control vs DREADD | n=7-8 | No | Mann-Whitney Test | **p= n.s** |
| | R. Distance Travelled female Control vs DREADD | n=7-8 | Yes | Unpaired t Test | **t=1,436, df=13, p=n.s** |
| | S. Velocity female Control vs DREADD | n=7-8 | Yes | Unpaired t Test | **t=1,436, df=13, p=n.s** |
